# TOR signaling modulates Cdk8-dependent *GAL* gene expression in *Saccharomyces cerevisiae*

**DOI:** 10.1101/2020.05.15.097576

**Authors:** Nicole Hawe, Konstantin Mestnikov, Riley Horvath, Mariam Eji-Lasisi, Cindy Lam, John Rohde, Ivan Sadowski

## Abstract

Cdk8 of the RNA Polymerase II mediator complex regulates genes by phosphorylating sequence specific transcription factors. Despite conserved importance for eukaryotic transcriptional regulation, the signals regulating Cdk8 are unknown. Full induction of the yeast *GAL* genes requires phosphorylation of Gal4 by Cdk8, and we exploited this requirement for growth of *gal3* yeast on galactose to identify mutants affecting Cdk8 activity. Several mutants from the screen produced defects in TOR signaling. A mutant designated <u>g</u>al <u>f</u>our throttle (*gft*) 1, *gft1*, was identified as an allele of *hom3*, encoding aspartokinase. Defects in *gft1*/ *hom3* caused hypersensitivity to rapamycin, and constitutive nuclear localization of Gat1. Furthermore, mutations of *tor1* or *tco89*, encoding TORC1 components, also prevented *GAL* expression in *gal3* yeast, and *tco89* was determined to be allelic to *gft7*. Disruption of *cdc55*, encoding a subunit of PP2A regulated by TOR signaling, suppressed the effect of *gft1*/ *hom3, gft7*/ *tco89*, and *tor1* mutations on *GAL* expression in *gal3* yeast, but not of *cdk8*/ *srb10* disruptions or Gal4 S699A mutation. Mutations of *gft1*/ *hom3* and *tor1* did not affect kinase activity of Cdk8 *in vitro*, but caused loss of Gal4 phosphorylation *in vivo*. These observations demonstrate that TOR signaling regulates *GAL* induction through the activity of PP2A/ Cdc55, and are consistent with the contention that Cdk8-dependent phosphorylation of Gal4 S699 is opposed by PP2A/ Cdc55 dephosphorylation. These results provide insight into how induction of transcription by a specific inducer can be modulated by global nutritional signals through regulation of Cdk8-dependent phosphorylation.

## Introduction

Cdk8 is a protein kinase of the eukaryotic RNA Polymerase II mediator complex, that modulates gene expression in response to sequence-specific transcription factors. Phosphorylation of transcriptional activators by Cdk8 produces positive or negative effects on gene expression, depending on the functional effect of the modification (1). The duplicitous effect of Cdk8 on gene regulation is conserved in humans, but is best understood in yeast, where mutants of *cdk8* were identified in numerous screens for alterations in gene regulation, which resulted in a proportion of aliases that reflects its diverse effect on gene regulation, including *ssn3, srb10, ume5, gig2, nut7, ssx7, urr1*, and *rye5* (2-5). Cdk8 activity is dependent upon additional proteins of the mediator kinase module, including cyclin C/ Srb11, Srb8/ Med12 and Srb9/ Med13 (6). Several observations indicate that Cdk8 activity in yeast is regulated by nutrient availability. Cdk8 is degraded in nitrogen starved cells (7), and its abundance is reduced as nutrients become depleted (3). Additionally, the associated cyclin C/ Srb11 is degraded in response to oxidative stress and carbon limitation (8, 9). Remarkably, despite its role in regulating responses to multiple physiological signals in yeast, a function that is likely conserved in humans (7), signaling mechanisms that regulate Cdk8 activity and phosphorylation of its substrates have not been identified.

Several yeast Cdk8 transcription factor substrates, including Ste12 and Phd1 (7, 10), regulate filamentous growth (FG) in response to nutrient limitation, where cells differentiate into elongated filamentous forms to promote nutrient foraging (11). Nitrogen limitation inhibits Cdk8 activity (7), which allows accumulation of these factors to cause induction of genes that promote FG. In contrast, other transcriptional activators, notably Gal4, are positively regulated by Cdk8 phosphorylation (12), where phosphorylation is required for full induction of the *GAL* genes in response to galactose (13). Additional transcriptional activators that are positively regulated by Cdk8 include Sip4 (14) and Skn7 (15). For each of these regulatory effects, activity of Cdk8 is associated with a favorable growth environment, and Cdk8 activity and phosphorylation of its substrates are inhibited under conditions of nutrient or physiological stress (7, 8,13, 15, 16), although as mentioned above, signaling mechanisms that regulate Cdk8 activity have not been identified.

Yeast respond to their nutritional environment through multiple signaling mechanisms that include the RAS-cAMP-PKA, SNF1/ AMPK, and target of rapamycin (TOR) pathways, which are interconnected by cross-talk, and are conserved amongst eukaryotes (17-20). TOR signaling regulates cell growth by promoting macromolecular synthesis in response to nutrients, particularly amino acids, while inhibiting catabolic processes and autophagy (21). TOR-complex 1 (TORC1), comprised of the partially redundant Tor1 or Tor2 protein kinases, and the regulatory subunits Lst8, Kog1 and Tco89, responds to the abundance and quality of nutrients, primarily nitrogen (22), and is inhibited by the fungal antibiotic rapamycin in a complex with the prolyl isomerase FKBP12/ Fpr1, to produce an effect that mimics nutrient starvation in yeast and human cells (19). TORC1 activity stimulates protein translation, ribosome biogenesis, cell cycle progression, and regulates nutrient uptake and amino acid metabolism (22). These effects are mediated by multiple downstream targets, including Sch9 (23), the yeast homologue of ribosomal protein S6 kinase, and Tap42, which inhibits the activity of two protein phosphatase complexes represented by the type 2A phosphatase Pph21/ Pph22/ Pph3, and 2A-related phosphatase Sit4 (18). A significant proportion of transcriptional responses to nutrients regulated by TORC1 involves dephosphorylation of transcription factors by these phosphatases. For example, inhibition of TORC1 by nitrogen limitation causes activation of PP2A and Sit4 which dephosphorylate the transcription factors Gat1 and Gln3, allowing translocation to the nucleus for activation of genes that are normally repressed under ideal growth conditions, an effect known as nitrogen catabolite repression (NCR) (24,25). Most studies of transcriptional responses involving TOR have focused on nitrogen availability, but TORC1 activity is also inhibited by additional stress conditions, including glucose limitation, an effect that requires AMPK/ Snf1 (19, 26). Glucose starvation activates Snf1/ AMPK kinase activity, which inhibits TORC1 by a mechanism involving formation of vacuole-associated Kog1 bodies, which prevents phosphorylation of Sch9 (27). However, overall the mechanism(s) by which glucose and carbon signaling modifies activity of transcription factors regulated downstream of TORC1 have not been characterized.

The galactose-induced genes (*GAL*) in yeast have served as an important model for understanding mechanisms of transcriptional regulation in eukaryotes (28). The *GAL* genes are activated by Gal4, which in the absence of galactose is inhibited by direct interaction with Gal80 (29). Galactose induces *GAL* expression as a ligand of Gal3 protein, which relieves the inhibitory effect of Gal80 on Gal4 (30, 31). Yeast with *gal3* mutations produce an interesting phenotype where *GAL* expression is induced several days post exposure to galactose, instead of hours in WT yeast, an effect termed long term adaptation (LTA) to galactose (32). Characterization of the *gal3* LTA phenotype revealed that full *GAL* induction requires phosphorylation of Gal4 by Cdk8 at S699, and consequently that *GAL* gene expression is modulated by signals that control Cdk8 (12, 13), although as mentioned, signals controlling Cdk8 or phosphorylation of its substrates have not been identified.

To identify mechanisms regulating Cdk8, we conducted a genetic screen which exploited the dependence of Cdk8 for growth of *gal3* yeast on galactose. From this effort we identified mutants that prevent *GAL* expression only in combination with a *gal3* null allele, designated the <u>g</u>al <u>f</u>our throttle mutants (*gft*). Characterization of one class of this mutant collection revealed that phosphorylation of Gal4 by Cdk8 is opposed by PP2A/ Cdc55 phosphatase downstream of TORC1. Collectively, we demonstrate here that *GAL* induction in yeast is modulated by the additional nutrient environment, through a mechanism involving TOR signaling and a balance of phosphorylation of the regulatory Cdk8-dependent site on Gal4.

## Results

### A minor subset of *gal3* yeast induce full *GAL* expression in response to galactose

Yeast defective for *gal3* produce a distinctive long term adaptation (LTA) phenotype where induction of *GAL* expression occurs days post exposure to galactose rather than minutes to hours in wild type yeast (32). In a wild type yeast strain (W303) bearing a GFP reporter gene expressed from the *GAL1* promoter we find that ∼80% of cells induce significant GFP expression within an hour of galactose addition, and greater than 95% become fully induced within 4 hours (Figure 1A). In contrast, consistent with previous observations (33), only a minor subset of *gal3* yeast induce GFP expression even at 24 hours, and after 96 hours only ∼10% of the cells induce significant expression (Figure 1B). Importantly however, individual *gal3* cells are capable of inducing GFP expression comparable to levels similar to wild type. Robust induction of *GAL* expression in a minor proportion of *gal3* yeast is also observed by growth on plates containing ethidium bromide (EB), with galactose as the sole carbon source (EB-gal). EB inhibits mitochondrial function and forces utilization of sugars as carbon source (33); consequently, growth on EB-gal is a stringent measurement of *GAL* gene induction. Consistent with results using the GFP reporter, we find that approximately 10% of *gal3* yeast in the W303 background produce colonies on EB-gal plates of similar size as wild type (Figure 2). Importantly, disruption of *cdk8* or mutation of the Cdk8-dependent phosphorylation site on Gal4 at S699 completely prevents growth on EB-gal plates (13).

**Figure 1:**
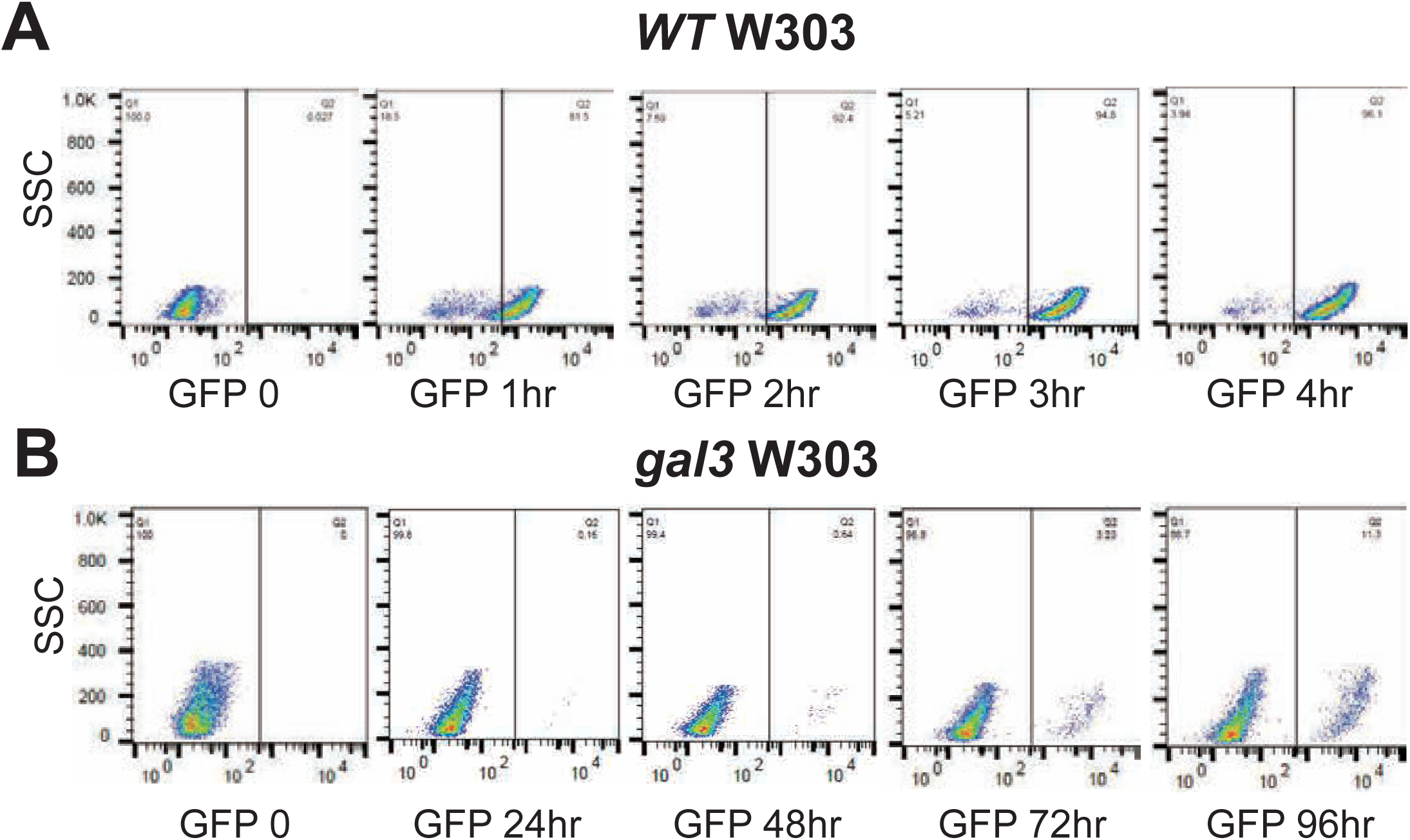
Yeast defective for *gal3* produce delayed induction of a *GAL-GFP* reporter gene. Wild type W303-1A yeast (yNH029) (**Panel A**) or *gal3* (yNH30) (**Panel B**) were grown in minimal media containing glycerol, lactic acid and ethanol (GFP 0), and induced by addition of galactose to 2% for the indicated time (hrs) when samples were taken for analysis by flow cytometry. GFP fluorescence is indicated on the X-axis and side scatter on the Y-axis.

**Figure 2:**
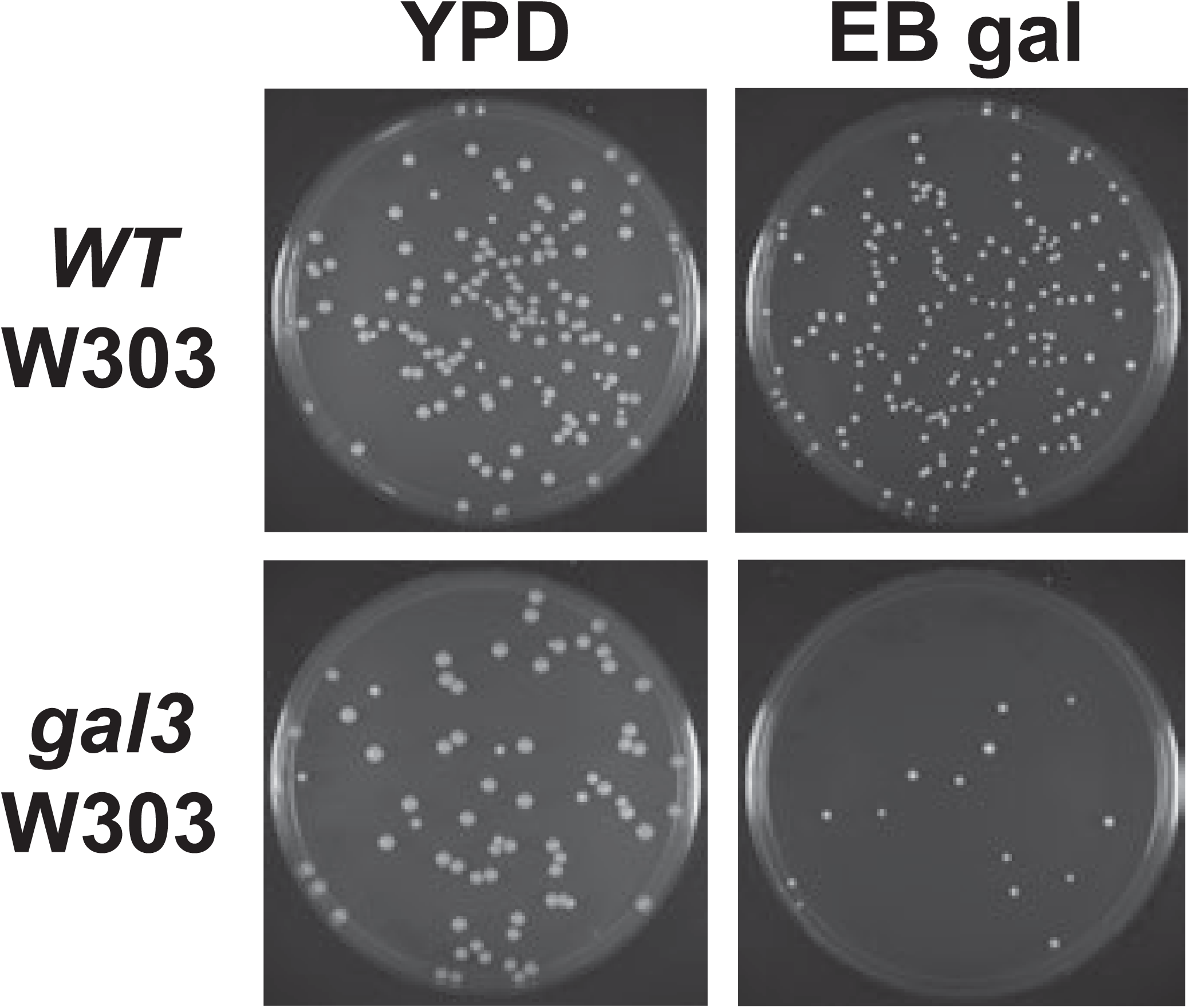
Yeast defective for *gal3* produce rare but robust colonies on EB-gal plates, typical of the long-term adaptation phenotype. WT and *gal3* (ISY54) yeast were grown overnight in YPD, diluted to equivalent O.D. A_600_, plated on YPD (left) or EB-gal plates (right), and incubated at 30°C for 5 days.

### *Hom3*/ *gft1* is necessary for long term adaptation/ delayed induction of *gal3* yeast

To identify factors that control Cdk8 activity and phosphorylation of Gal4 S699 we used a genetic screen to isolate mutants of genes necessary for growth of *gal3* yeast on EB-gal. Because less than 10% of otherwise wild type *gal3* yeast produce a colony on EB-gal (Figure 2), we were unable to implement a conventional synthetic genetic array (SGA) screen using the non-essential gene deletion collection for this purpose. Consequently, we employed u.v.-irradiation of a *gal3* W303 strain and replica plating, to identify mutants that were incapable of growth on EB-gal, but which produced robust colonies on media containing 3-carbon molecules as the sole carbon source. Initial mutants recovered from this process were transformed with a plasmid bearing genomic *GAL3* and re-assayed for growth on EB-gal; mutants capable of growth when expressing Gal3 were designated the <u>g</u>al <u>f</u>our throttle (*gft*) mutants, a collection which is comprised of several groups with distinct phenotypes that will be detailed in a separate report. The identity of one mutant, *gft1*, was identified by complementation with a plasmid library from WT yeast, as a recessive allele of *hom3*, which encodes aspartate kinase. Transformation with plasmids bearing a genomic fragment of *HOM3*, or expressing the Hom3 ORF from the *TEF1* promoter, allow growth of the *gal3 gft1* strain on EB-gal plates (Figure S1). Additionally, disruption of *hom3* in a W303 *gal3* strain prevents growth on EB-gal (Figure 3A), and also inhibits LTA to galactose as measured using a *GAL1*-GFP reporter (Figure 3B), or *GAL1*-LacZ reporter (Figure S2). Furthermore, a *hom3* null allele was non-complementing with the *gft1* mutation (not shown), and we found that *gft1-1* mutant strains do not produce Hom3 protein as determined by immunoblotting (Figure 4B, lane 5).

**Figure 3:**
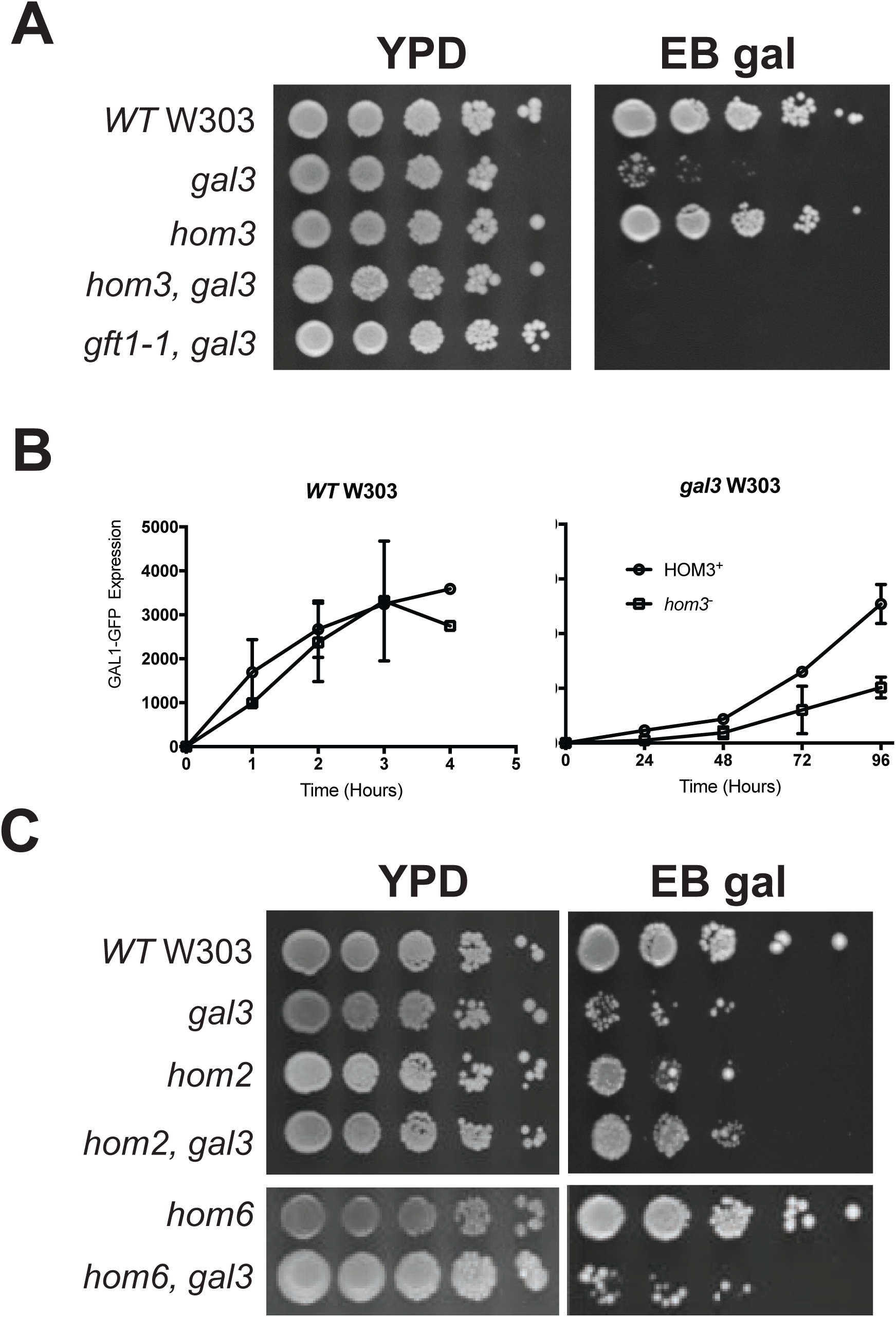
Aspartate kinase/ Hom3 is required long term adaptation. **Panel A:** Strains with the indicated genotype were grown overnight in YPD, diluted to an O.D. A_600_ of 1.0, spotted onto YPD or EB-gal plates in 10-fold serial dilutions, and grown at 30°C for 5 days. **Panel B:** WT (left) or *gal3* (right) yeast bearing WT *HOM3* (yNH029, yNH030) (○) or a *hom3* disruption (yNH031, yNH032) (□) were induced with 2% galactose for the indicated time (hrs) and analyzed by flow cytometry. Results are presented as mean fluorescence intensity (MFI) of GFP expression and represent an average from three independent cultures. **Panel C:** Strains with the indicated genotype were diluted, spotted onto YPD or EB-gal plates in 10-fold serial dilution and grown at 30°C for 5 days.

**Figure 4:**
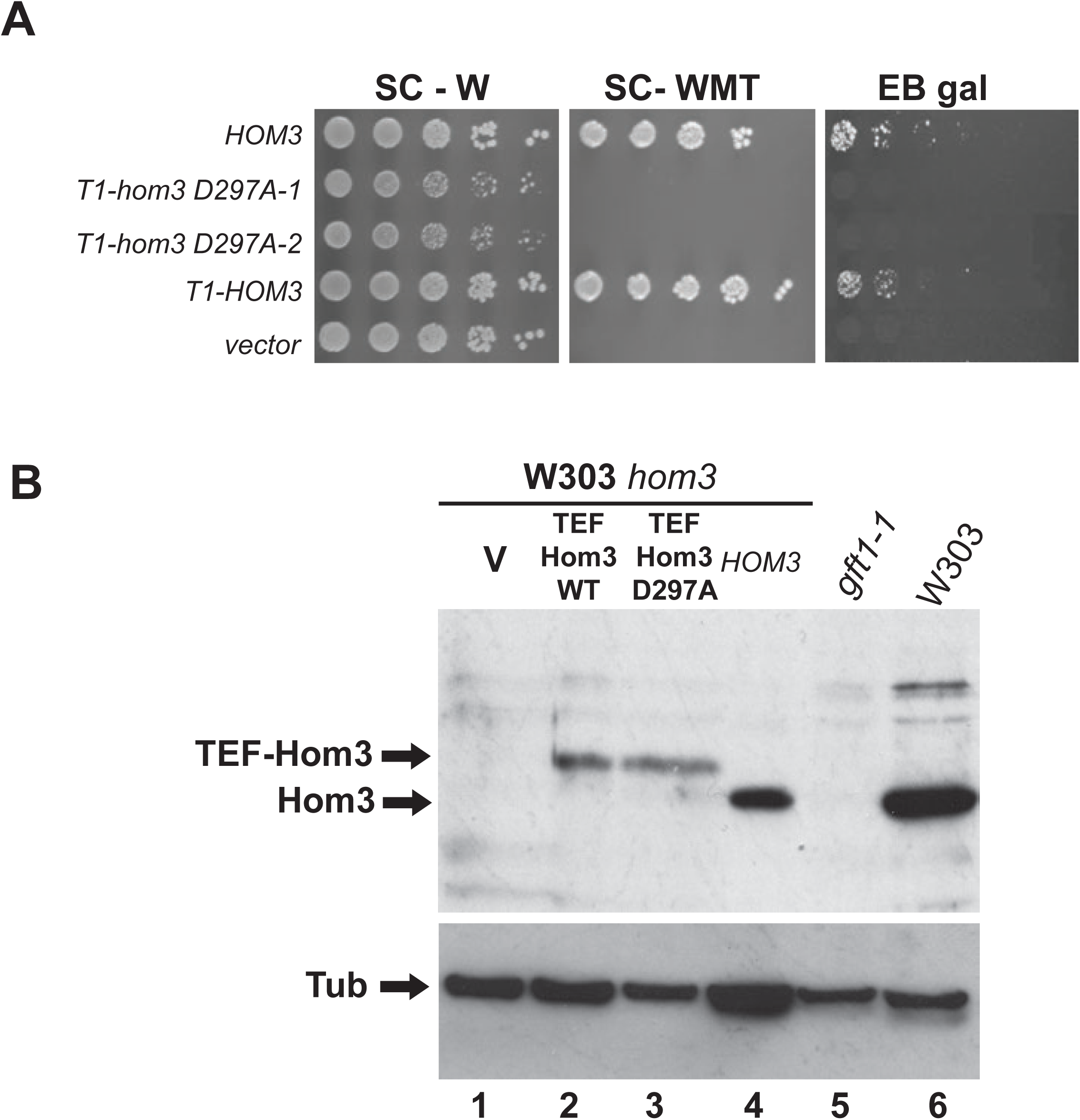
Aspartate kinase/ Hom3 catalytic activity is required for long term adaptation. **Panel A:** A *gal3 hom3* strain (ISY128) was transformed with plasmids bearing a genomic clone of *HOM3* (pIS297), a vector control (pRS314), T1-*HOM3* (pIS574) expressing the Hom3 ORF from the *TEF1* promoter, or two clones of pNH01 expressing the Hom3 D297A mutation. Cultures were diluted and spotted onto SC lacking tryptophan (SC-W), SC lacking tryptophan, methionine and threonine (SC-WMT), or EB-gal, from 10-fold serial dilutions. **Panel B:** Protein extracts were prepared from WT W303-1A (lane 6), *gal3 gft1-1* (ISY135, lane5), or *hom3* (ISY279, lanes 1-4) strains transformed with plasmids expressing Hom3 from a genomic clone (pIS297, lane 4), Hom3 WT (pIS574, lane 2), the D297A mutant (pNH01, lane3) from the *TEF1* promoter, or a vector control (pRS314, lane 1). Samples were resolved on SDS-PAGE and analyzed by immunoblotting with antibodies against Hom3 (top panel) or tubulin (bottom). Arrows indicate migration of specific proteins produced from the TEF1-Hom3 and genomic Hom3 plasmids (top).

### Long term adaptation to galactose requires Hom3 catalytic activity but not homoserine biosynthensis

Aspartate kinase (Hom3) is the first enzyme in the pathway for synthesis of homoserine, the precursor for threonine and methionine biosynthesis (Figure S3). We examined whether a general defect in this pathway may prevent LTA of *gal3* yeast, by examining the effect of *hom2* and *hom6* disruptions, which affect aspartic beta semi-aldehyde dehydrogenase, and homoserine dehydrogenase, respectively downstream of *HOM3* (Figure S3). Neither of these disruptions affected growth on EB-gal plates (Figure 3C), in either WT or *gal3* W303 yeast, indicating that a defect in *hom3* specifically impairs LTA in *gal3* yeast, rather than a general deficiency in this metabolic pathway. Additionally, we found that mutation of Hom3 aspartate 297 to alanine (D297A), a residue conserved within the catalytic domain of kinase enzymes (34), causes threonine and methionine auxotrophy, and also prevents growth of *gal3* yeast on EB-gal plates (Figure 4A). The D297A *hom3* mutation does not affect abundance of Hom3 protein (Figure 4B), which indicates that Hom3/ aspartate kinase catalytic activity is required for LTA and not merely the protein itself.

### Mutation of hom3 causes a defect in TOR signaling

The Hom3 protein was previously shown to interact with the peptidyl-prolyl isomerase FKBP12/ Fpr1, and this interaction is required for feedback inhibition of aspartate kinase activity by threonine (35, 36). FKBP12 also binds the immunosuppressive agent rapamycin, and this receptor ligand complex inhibits TOR kinase function by direct interaction (37). Relating to these observations, we found that strains bearing the *gft1-1* mutation (not shown), and *hom3* disruption strains, were more sensitive to sub-lethal concentrations of rapamycin compared to wild type W303 or a strain bearing a *gal3* disruption (Figure S4A). This observation is consistent with previous analysis indicating that *hom3* mutants are amongst the most sensitive to rapamycin within the yeast haploid deletion set (38), and sensitivity to sub-lethal concentrations of rapamycin is known to indicate defects in TOR pathway signaling (39). Because of the previously described relationship between Hom3 and FKBP12 we examined whether this protein may also affect *GAL* expression, but found that disruption of *fpr1* did not affect growth of *gal3* W303 yeast on EB-gal, nor did it suppress the effect of the *hom3* disruption on the *gft* phenotype (Figure S4B).

Multiple nutrient responsive transcription factors, including the GATA factors Gat1 and Gln3 are negatively regulated by TOR signaling, where nutrient limitation, or treatment with rapamycin, causes their dephosphorylation allowing translocation to the nucleus for regulation of target genes (24, 25). We examined the effect of *hom3* null mutations on subcellular localization of Gat1 using a GFP fusion, where we found it to be constitutively localized to the nucleus in cells growing in rich medium, similar to yeast bearing a disruption of *tor1*, and in contrast to wild type yeast where the fusion is predominately cytoplasmic (Figure 5). These results suggest that defects in aspartate kinase cause inhibition of TOR signaling.

**Figure 5:**
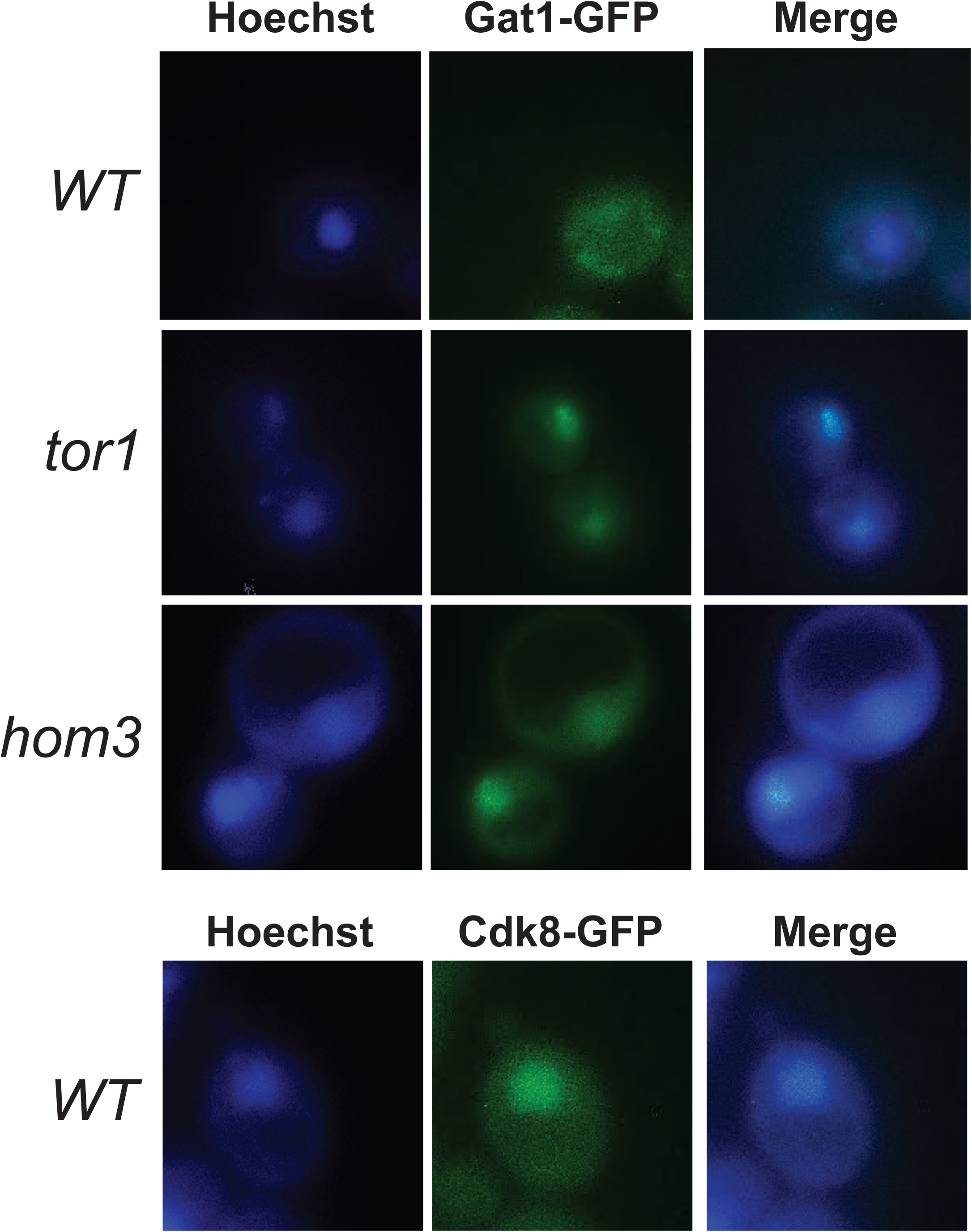
Defects in aspartate kinase/ Hom3 causes constitutive nuclear localization of Gat1. Strains W303-1A (WT), *tor1* (yNH008), and *hom3* (ISY279) expressing Gat1-GFP (pRS314-GAT-GFP, top panels), or Cdk8-GFP (pJP015, bottom panel) were examined by fluorescent microscopy. DNA stain (Hoechst), GFP, and merged images are shown from representative samples.

TORC1 protein kinase activity can be measured *in vitro* using a semi-intact cell assay system, where activity is measured by phosphorylation of the translation regulatory factor 4-EBP1 (40). We used this assay to examine whether *hom3* mutation causes a defect in TORC1 kinase function. Consistent with the initial report describing this assay, we observed phosphorylation of 4-EBP1 in reactions with preparations from untreated wild type cells (Figure 6, lane 1), which was significantly reduced in reactions treated with 10 μM rapamycin (lane 5), which confirms that the assay can detect inhibitory effects on TORC1 activity. However, we found that TORC1 activity in this assay was unaffected in preparations from *gal3* or *hom3* strains (lanes 2 and 3), indicating that either the *hom3* mutation must inhibit signaling downstream of TORC1, or that an *in vivo* inhibitory effect is abrogated during preparation of the samples. A further caveat is that the TORC1 complex can utilize either Tor1 or Tor2 (22), and consequently it is possible that the effect of *hom3* deletions on Tor1 activity may be obscured by redundancy for phosphorylation of 4EBP1 in this assay.

**Figure 6:**
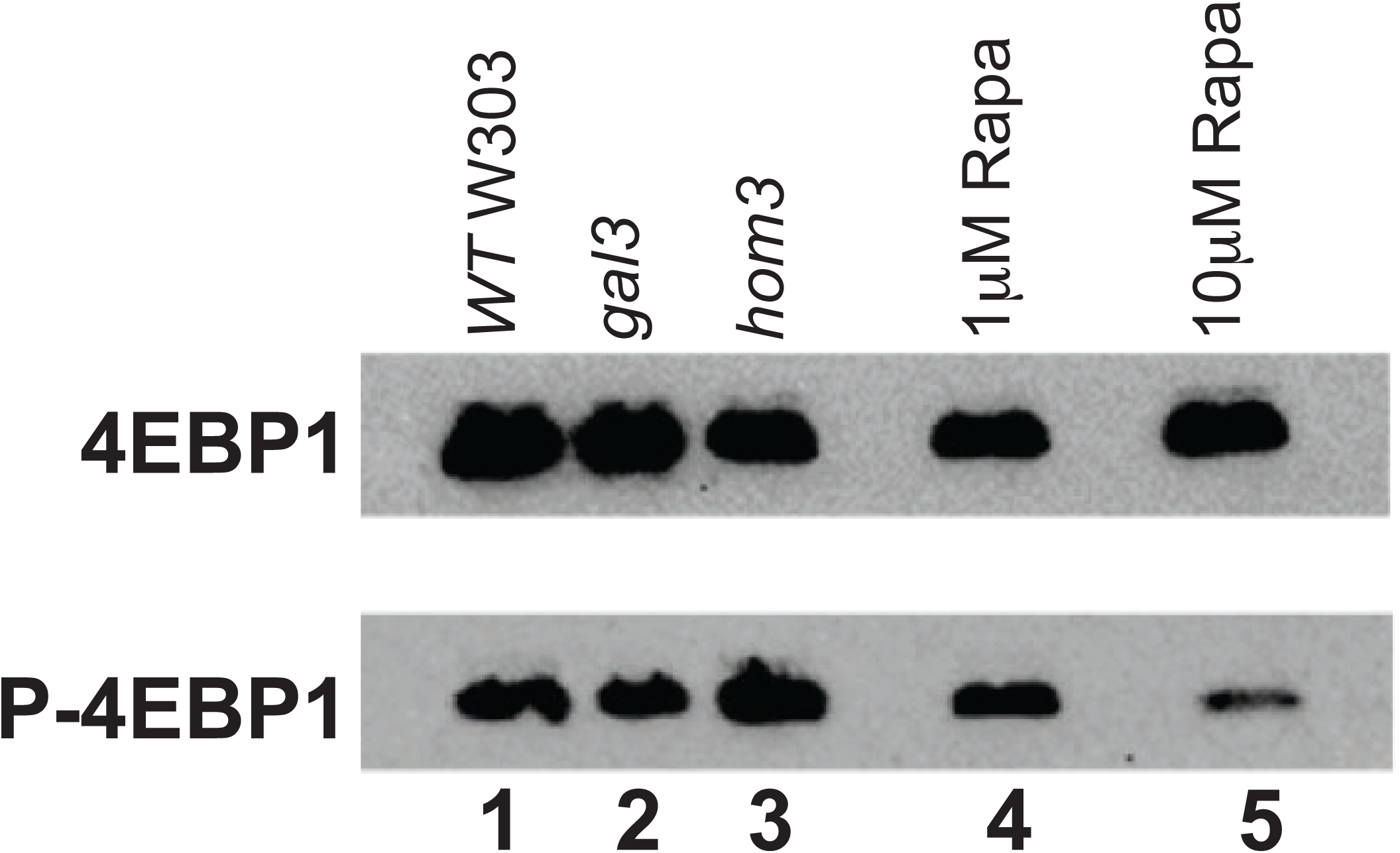
Deletion of *hom3* does not affect TORC1 kinase activity *in vitro*. TORC1 activity was assayed in semi-permeable cell samples prepared from WT (lanes 1, 4 and 5), *gal3* (ISY54, lane 2), and *hom3* (ISY279, lane 3). Reactions in lanes 4 and 5 contained 1 or 10 µM rapamycin, respectively. Samples were analyzed by immunoblotting for total 4EBP1 (bottom panel) and phosphorylated 4EBP1 (top panel).

### Defects in TOR signaling prevent long term adaptation to galactose

Because *hom3* strains were found to be hypersensitive to sublethal concentrations of rapamycin, we examined whether additional *gft* mutants produced this phenotype, and found that 5 further mutants from the screen were more sensitive to rapamycin than the parental W303 strain, although amongst these the *gft1*/ *hom3* mutant was the most sensitive (not shown). We then examined whether disruption of genes encoding components of TOR signaling also produce the *gft* phenotype in combination with a *gal3* mutation. The only non-essential genes encoding proteins of the TORC1 complex are *TOR1*, and *TCO89* (22). Interestingly, we found that disruption of *tco89* in a *gal3* W303 strain, prevents growth on EB-Gal plates, but disruption in a WT strain has no effect (Figure 7A). Consistent with the growth assays on EB-Gal, disruption of *tco89* on its own did not affect induction of a *GAL*-GFP reporter gene (Figure 7B, left panel), but prevents induction in combination with a *gal3* disruption (Figure 7B, right), indicating that *tco89* disruption prevents LTA, equivalent to the effect of the *gft* mutants. Consequently, we examined whether mutants from the *gft* screen might represent *tco89* alleles, where we found that *tco89* disruptions were non-complementing with the mutant we had designated *gft7*; diploid cells produced by mating *gft7* and *tco89* haploids did not produce spores capable of growth on EB-gal, confirming that the *gft7* mutation is allelic to *tco89* (Figure S5)

**Figure 7:**
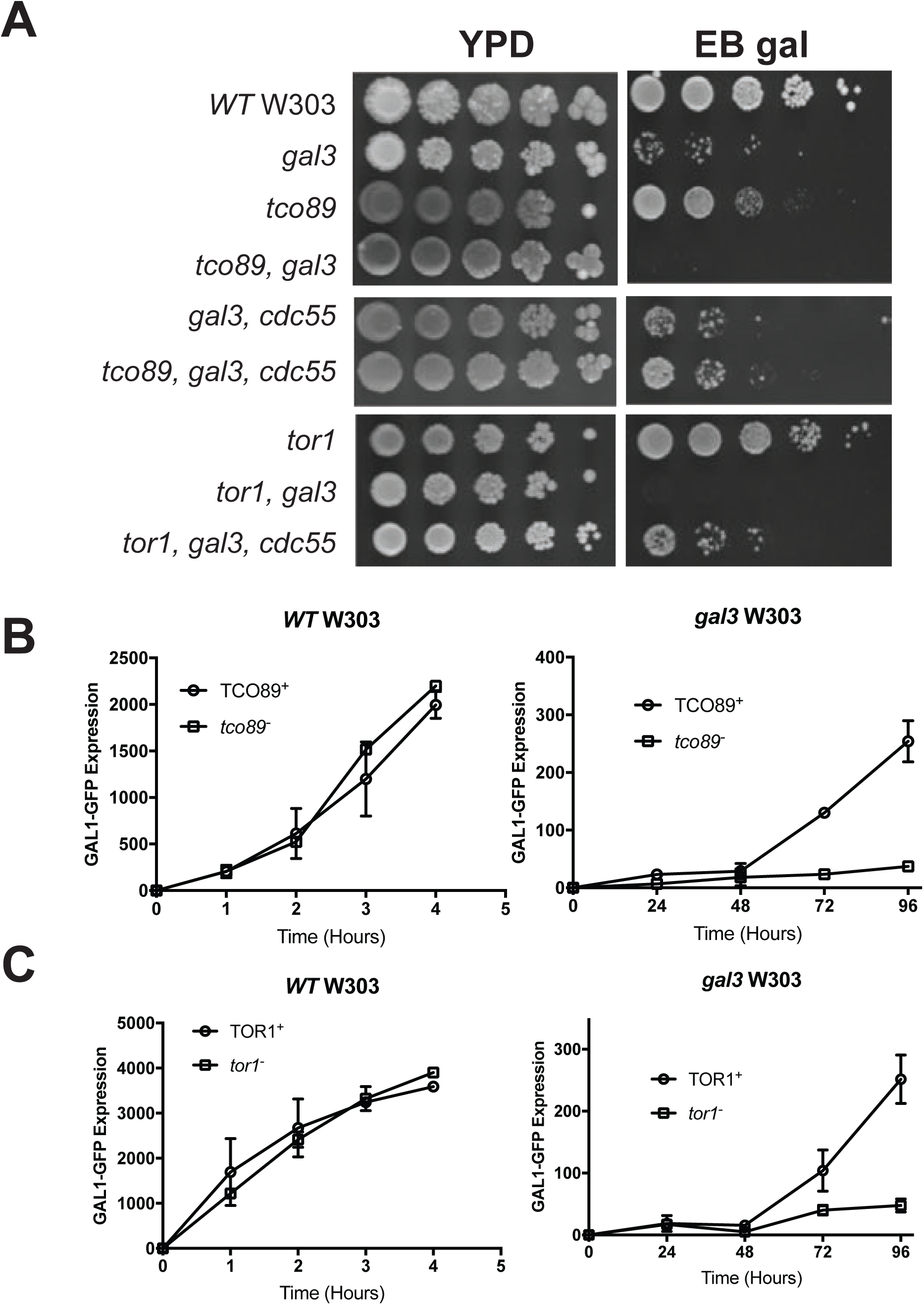
Mutation of TORC1 subunits prevents LTA in *gal3* yeast. **Panel A:** Strains with the indicated genotype were diluted, spotted onto YPD or EB-gal plates in 10-fold serial dilutions, and grown at 30°C for 5 days. **Panel B:** WT (left panel) or *gal3* (right) yeast bearing WT *TCO89* (yNH029, yNH030) (○) or a *tco89* disruption (yNH035, yNH036) (□) were induced with 2% galactose for the indicated time (hrs) and analyzed by flow cytometry. Results are presented as mean fluorescence intensity (MFI) of GFP expression and represent an average from three independent cultures. **Panel C:** WT (left panel) or *gal3* W303-1A (right) yeast bearing WT *TOR1* (yNH029, yNH030) (○) or a *tor1* disruption (yNH033, yNH034) (□) were induced with 2% galactose for the indicated time (hrs) and analyzed by flow cytometry. Results are presented as mean fluorescence intensity (MFI) of GFP expression and represent an average from three independent cultures.

*TOR1* is non-essential for yeast growth, and redundant with *TOR2* for TORC1 activity (22). We found that disruption of *tor1* in WT yeast did not affect on growth on EB-gal, but completely prevented growth of *gal3* strains (Figure 7A), and thus also produces the *gft* phenotype, similar to the effect of *hom3* and *tco89* mutants. Disruption of *tor1* also inhibits induction of *GAL*-GFP expression in a *gal3* strain (Figure 7C, right panel), but not in WT cells (left). However, unlike *tco89*, in complementation analysis we found that *tor1* disruption was not allelic to any of the *gft* mutants. Rather, *tor1* disruption produced *gft* mutant phenotype:WT spores at a 3:1 ratio from segregants of crosses with the *gft2, gft6* and *gft14* mutants (Figure S6), representing possible unlinked non-complementation. This may indicate that these additional 3 mutants may also cause defects in TOR signaling. Interestingly, we note that non-allelic non-complementation was observed while characterizing the original *tor* mutants (37). Taken together, these observations demonstrate that induction of the *GAL* genes by LTA in response to galactose requires TORC1, and specifically the Tor1 protein kinase.

### PP2A-Cdc55 phosphatase inhibits GAL induction

TOR signaling is known to inhibit several downstream protein phosphatases which control nutrient-responsive gene expression (22). Sit4 is a type 2A-related protein phosphatase regulated by TOR, and known to dephosphorylate multiple nutrient-responsive transcription factors, including Gat1 and Gln3 (41) (24). We examined whether Sit4 might modulate *GAL* induction, and found that *sit4* disruption did not affect growth of wild type or *gal3* yeast strains on EB-Gal (Figure S7), indicating that Sit4 is not required for *GAL* induction. Furthermore, *sit4* disruption also did not allow growth of *hom3 gal3* yeast on EB-Gal, which suggests that Gal4 activity is likely not affected by Sit4 phosphatase (Figure S7).

TOR signaling also regulates protein phosphatase 2A (PP2A) comprised of the redundant Pph21/ Pph22 catalytic subunits and the non-essential Tpd3 and Cdc55 regulatory subunits (42). To examine if PP2A regulates *GAL* induction, we determined the effect of *cdc55* disruptions on growth of W303 strains on EB-gal. Here we found that *cdc55* disruption had no effect on wild type or *gal3* W303, but interestingly suppressed the *gft* phenotype of *gal3 hom3* mutants (Figure 8A). Similarly, we found that *cdc55* disruption also suppressed the effect of *tor1* and *tco89* mutations for growth of *gal3* yeast on EB-gal (Figure 7A), indicating that defects in TOR signaling for *GAL* expression must involve PP2A/ Cdc55, which suggests that PP2A might counteract the effect of Cdk8 on Gal4.

**Figure 8:**
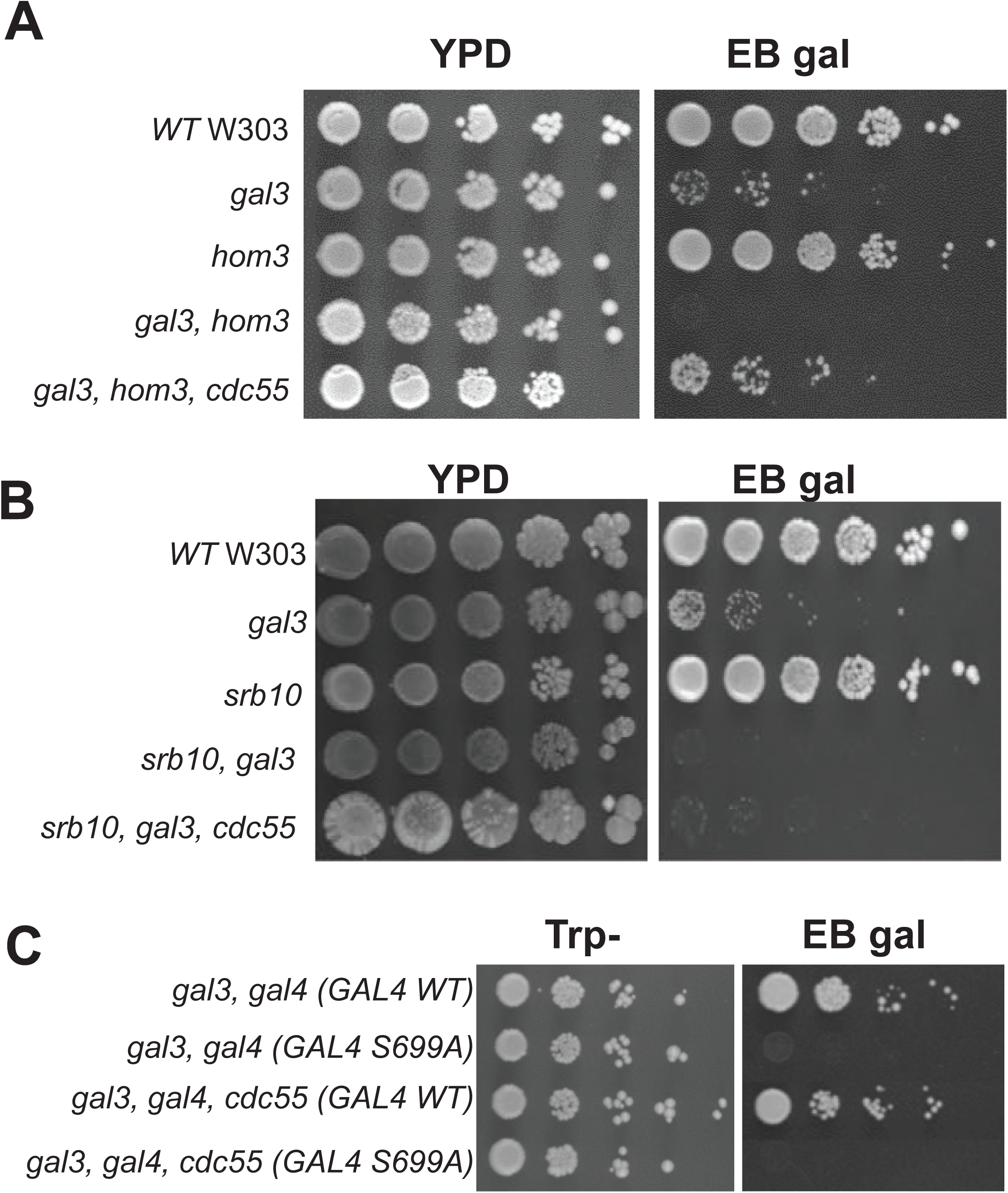
Disruption of *cdc55* suppresses effects of *hom3, tor1* and *tco89* on LTA in *gal3* yeast. **Panels A and B:** Strains with the indicated genotype were grown overnight in YPD, diluted to an O.D. A_600_ of 1.0, spotted onto YPD or EB-gal plates in 10-fold serial dilutions, and grown at 30°C for 5 days. **Panel C:** Strains *gal3 gal4* (YJR40) and *gal3 gal4 cdc55* (yNH037) were transformed with plasmids expressing WT GAL4 (YCpG4trp) or the GAL4 S699A mutant (pRD038), and cells spotted from liquid SC-Trp cultures in 10-fold serial dilutions on SC-Trp or EB gal plates, and grown for 5 days at 30°C.

Cdk8-dependent induction of *GAL* gene expression requires phosphorylation of Gal4 at serine 699 (12, 13, 16). Accordingly, disruption of *srb10*/ *cdk8* has no effect on growth of WT yeast, but prevents growth of *gal3* yeast on EB-gal (Figure 8B), which is typical of the *gft* phenotype described above. However, unlike *hom3, tor1* and *tco89* mutants that affect TOR signaling, disruption of *cdc55* does not suppress the effect of *srb10*/ *cdk8* mutants for growth of *gal3* yeast on EB-gal (Figure 8B). Similarly, mutation of the Cdk8 phosphorylation site on Gal4 at S699 to alanine also prevents growth of *gal3* yeast on EB-gal, and this effect is also not suppressed by disruption of *cdc55* (Figure 8C). These observations are consistent with the hypothesis that PP2A/ Cdc55 must affect *GAL* induction by altering Cdk8-dependent phosphorylation of Gal4 at S699.

### Defects in Tor signaling inhibit Gal4 phosphorylation in vivo but not Cdk8 kinase activity

The *gft* genetic screen was devised to identify pathways required for Gal4 phosphorylation-dependent induction of *GAL* expression, and consequently, we expected that mutants would prevent phosphorylation at Gal4 S699 by causing inhibition of Cdk8 kinase activity, or enhancing dephosphorylation. To examine this, we transformed the mutant strains with a plasmid expressing Flag-epitope tagged Cdk8, and assayed kinase activity of complexes recovered by immunoprecipitation. In these assays Flag-Cdk8 from wild type cells phosphorylates GST-RNA PolII C-terminal domain (CTD) fusion protein substrate or recombinant Gal4 protein (Figure 9A, lanes 2 and 3), but not GST alone (lane 1). Flag-Cdk8 recovered from *gal3, hom3* or *tor1* strains also phosphorylated GST-CTD and Gal4 as efficiently as wild type (lanes 5-6, 8-9 and 11-12), which indicates that inhibition of TOR signaling does not cause direct effects on Cdk8 kinase activity. In contrast, we found that a separate class of the *gft* mutant collection caused either partial or complete inhibition of Flag-Cdk8 activity in this assay (in preparation).

**Figure 9:**
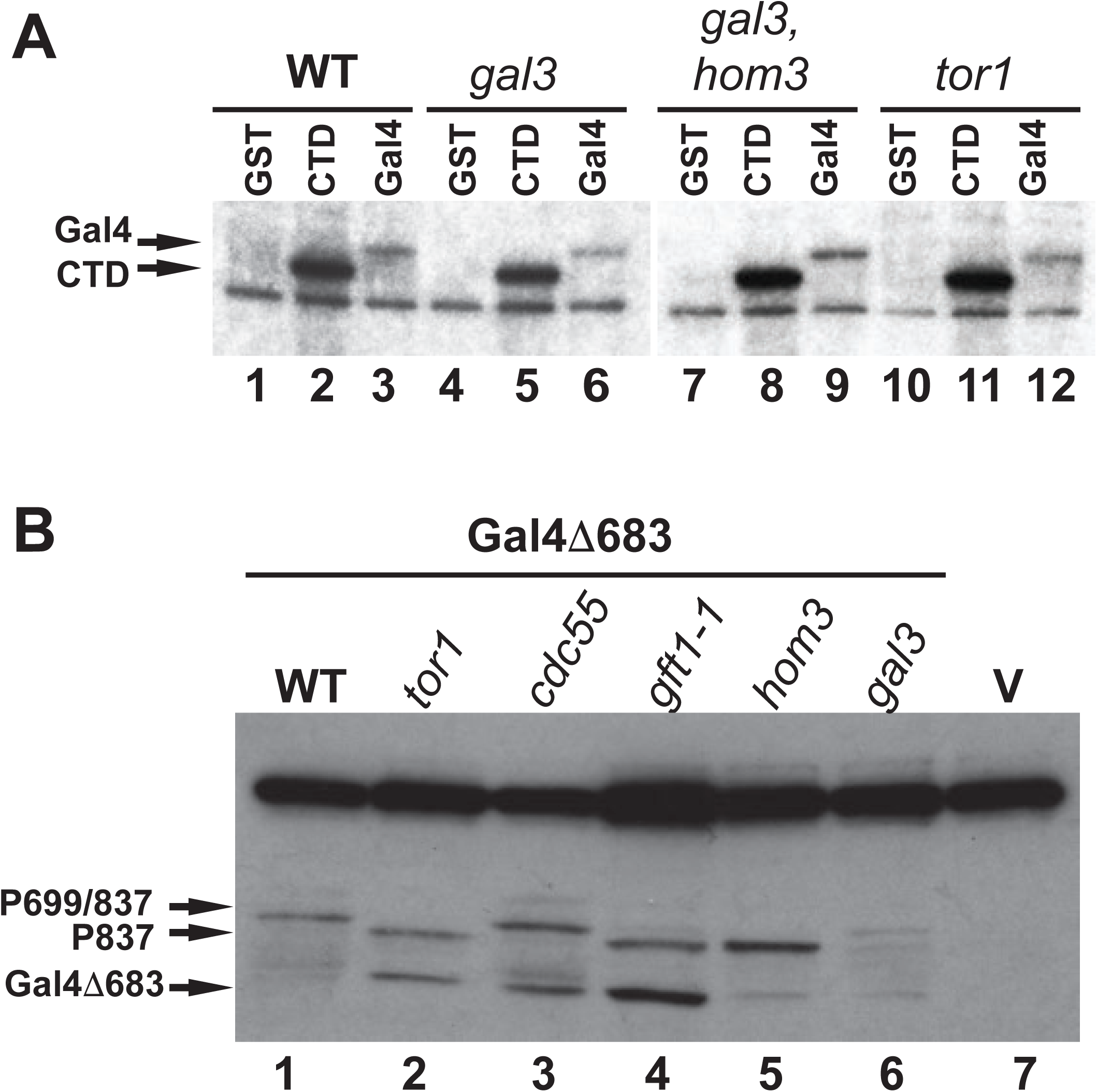
Mutations of *hom3* and *tor1* do not affect Cdk8 kinase activity, but alter Gal4 phosphorylation *in vivo*. **Panel A:** Cdk8-FLAG was recovered from strains W303-1A, *gal3* (ISY54), *gal3 hom3* (ISY128), and *tor1* (yNH008) by immunoprecipitation and used for *in vitro* kinase assays with GST (lanes 1, 4, 7, 10), GST fused to RNAPII CTD (lanes 2, 5, 8, 11) or recombinant Gal4 protein (lanes 3, 6, 9, 12) as the substrate. Reactions were analyzed by SDS-PAGE and autoradiography. **Panel B:** Protein extracts from *gal80* W303-1A strains (WT, lanes 1 and 7, YKM001), *tor1* (YKM003), *cdc55* (YKM002), *gft1-1* (YKM004), *hom3* (YKM005) and *gal3* (ISY54) expressing the Gal4Δ683 derivative (YCpG4trp, lanes 1-6) or a vector control (pRS314, lane 7) were analyzed by immunoblotting with antibodies against Gal4 DBD. Arrows indicate migration of unphosphorylated Gal4Δ683, and species produced by phosphorylation at S837 and S699 (P699/837) or S837 alone (P837).

We also examined the effect of *gft1*/ *hom3* and *tor1* mutations on Gal4 phosphorylation *in vivo*. Gal4 is phosphorylated on multiple sites which produce distinctive alterations in mobility in SDS PAGE (16). This effect is particularly exaggerated with the Gal4Δ683 mutant bearing a large central deletion, retaining only the N-terminal DNA binding domain (1-147) and C-terminal region including all of the verified *in vivo* sites of phosphorylation (683-881) (16). Additionally, because Gal4’s phosphorylations occur consequential to transcriptional activation (43) we expressed this protein in a *gal80* background where Gal4 activates transcription constitutively. The Gal4Δ683 protein produces several species detected by immunoblotting, representing the unphosphorylated and fully phosphorylated species which migrate at ∼40 and 46 KDa, respectively (Figure 9B, lane 1) in otherwise WT *gal80* W303. The slowest migrating Gal4 species is produced predominately by a combination of phosphorylations at S837 and S699 (16). Of these, only P-S699 is known to regulate Gal4 activity, which is produced by Cdk8 (12, 16). We also observe fully phosphorylated protein in strains with either *gal3* or *cdc55* disruptions (lanes 3 and 6). However, we note that strains bearing disruption of *tor1* (lane 2), *hom3* (lane 5), or the *gft1-1* mutation (lane 4) produce a faster migrating species typical of loss of the Cdk8-dependent P-S699 phosphorylation (16). These observations indicate that defects in Tor signaling inhibit phosphorylation of Gal4 at S699 *in vivo*.

Overall, these results presented here indicate that the *gft* genetic screen resulted in identification of genes required for Cdk8 activity, or that otherwise antagonize phosphorylation at Gal4 S699. For *gft1*/ *hom3*, this defect has revealed the role of TOR signaling, through PP2A/ Cdc55, for induction of *GAL* gene expression, which affects phosphorylation of Gal4 *in vivo* but not protein kinase activity of Cdk8. Taken together these observations suggest that PP2A/Cdc55 phosphatase may regulate *GAL* expression by dephosphorylation of Gal4 P-S699.

## Discussion

The yeast *GAL* genes represent an important model for understanding eukaryotic gene regulation in response to signal transduction. In this study we exploited the long-term adaptation (LTA)/ delayed *GAL* gene induction phenotype of *gal3* yeast, which is Cdk8-dependent (13), to identify regulators of Cdk8 using a mutant screen. One mutant, *gft1*, was identified as a recessive allele of *hom3*, which revealed a role for this enzyme in regulation of *GAL* expression but also a previously unrecognized effect of this enzyme on TOR signaling. Disruption of *hom3* prevents Cdk8-dependent induction of *GAL* expression in *gal3* yeast, and causes effects typical of defects in TOR signaling, including hypersensitivity to rapamycin and constitutive nuclear localization of the transcription factor Gat1 (24, 25). Furthermore, mutation of *tco89*, encoding a non-essential component of the TORC1 complex also prevented Cdk8-dependent *GAL* induction, and is allelic to *gft7*. Similar effects were caused by disruption of *tor1*, and genetic analysis indicated several additional mutants (*gft2, 6* and *14*) produce unlinked non-complementation with a *tor1* disruption, which suggests these may also encode proteins that regulate TORC1 (37). This indicates that at least 5 mutants from a screen to identify regulators of Cdk8-dependent *GAL* induction may produce defects in TOR signaling, which underscores the importance of this pathway for modulation of *GAL* gene expression.

Defects in TOR signaling caused by mutation of *hom3* or *tor1* inhibit induction of *GAL* expression in *gal3* yeast, but do not have direct effects on Cdk8 kinase activity. Rather, the original *gft1* mutation, as well as *hom3* and *tor1* disruptions inhibit production of the fully phosphorylated form of Gal4 *in vivo*, consistent with loss of the Cdk8-dependent phosphorylation at S699 (16). Effects of *hom3, tor1* and *tco89* on *GAL* expression in *gal3* yeast is suppressed by disruption of the protein phosphatase 2A (PP2A) regulatory subunit *cdc55*. These observations are consistent with known effects caused by inhibition of TORC1 activity, typified by dephosphorylation of transcription factors involved in nutrient response, including Gln3 and Gat1 which enables their nuclear translocation and activation of genes required for nutrient response (24, 25). Comparably, full induction of the *GAL* genes is dependent upon phosphorylation of Gal4 at S699 (12, 13), and consequently given our results, it seems likely that PP2A hyperactivation inhibits Cdk8-dependent *GAL* induction through dephosphoryation of Gal4 P-S699, which would implicate this specific phosphorylation as a regulatory substrate for PP2A (Figure 10).

**Figure 10:**
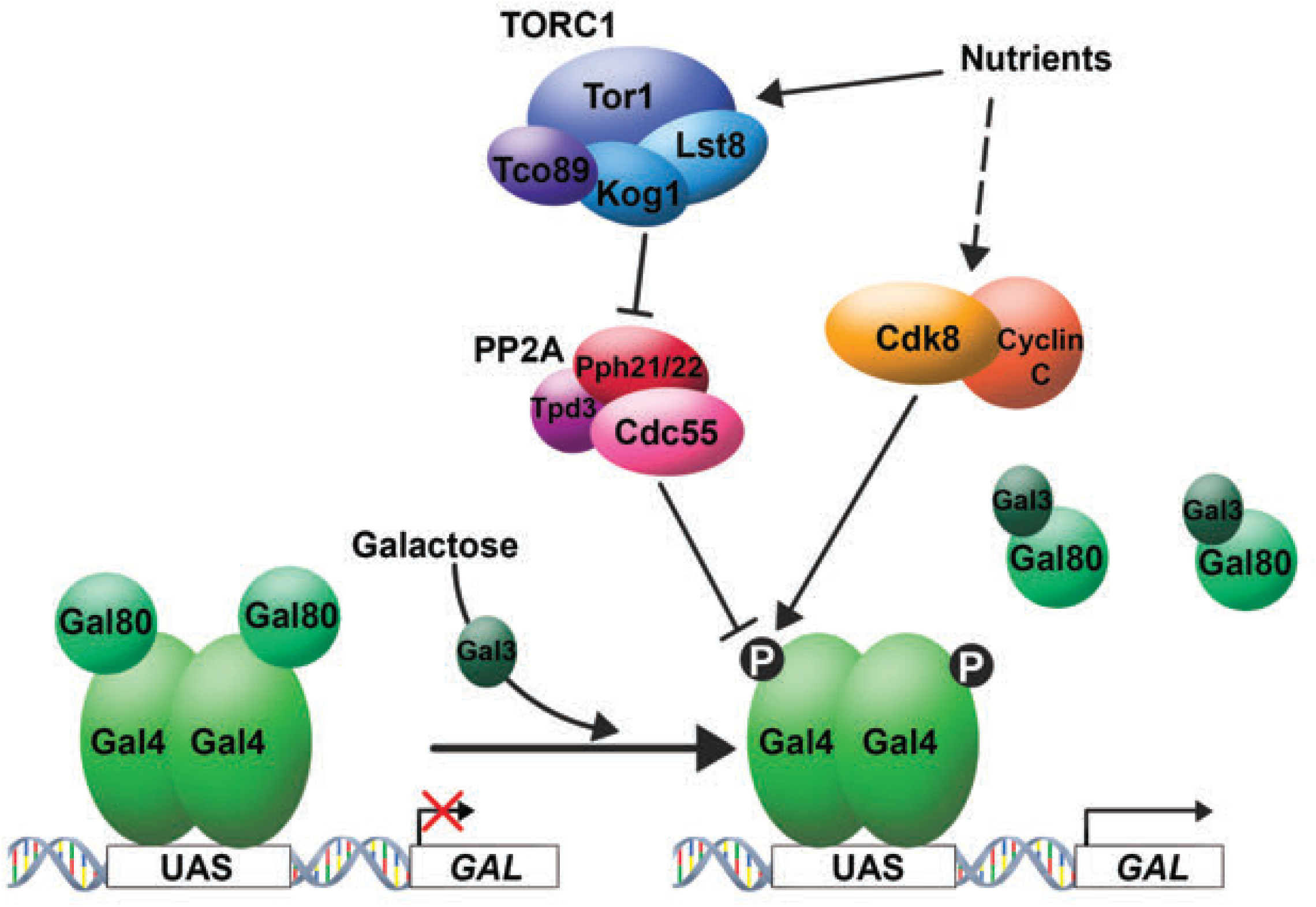
*GAL* induction is modulated by TOR signaling and PP2A/ Cdc55. Galactose induces *GAL* gene expression by binding the inducer protein Gal3, which relieves the inhibitory effect of Gal80 on the transactivator Gal4. Gal4 is phosphorylated by Cdk8 during transactivation on S699, which is required for full *GAL* gene induction. Nutrient signaling modulates *GAL* induction through TORC1 - PP2A signaling, and PP2A/ Cdc55 likely inhibits Gal4 activity by dephosphorylation of the Cdk8-dependent site at S699. Previous observations, and phenotypes produced by additional classes of mutants from the *gft* genetic screen indicate that Cdk8 activity itself is also regulated by nutrient signaling.

Regulation of TOR signaling is complex and involves sensing of nutrient availability through intracellular abundance of amino acids, most particularly glutamine (25). A relationship between aspartate kinase and TOR signaling was noted previously, in that *hom3* disruptions were found amongst the most sensitive to sub-lethal concentrations of rapamycin within the non-essential yeast deletion set (38). Furthermore, Hom3 physically interacts with the rapamycin receptor FKBP12/ Fpr1, which was shown to modulate feedback inhibition of Hom3 activity by threonine (35, 36). We found that *fpr1* disruption neither produced the *gft* phenotype in *gal3* yeast, nor did it suppress the effect of *hom3* mutations for *GAL* induction. Furthermore, mutations in *hom2* or *hom3*, encoding enzymes downstream of *HOM3* for synthesis of threonine and methionine (Figure S3) also had no effect on *GAL* induction in *gal3* yeast. The Hom3 D297A mutation produces the *gft* phenotype, indicating that specific loss of Hom3 catalytic activity, rather than general loss of this metabolic pathway causes inhibition of TOR signaling, although interestingly hyper-sensitivity to rapamycin appears to be typical of mutations of all gene components of this pathway (38). However, the fact that the *gft* phenotype is only produced by *hom3* mutation suggests that aspartate kinase has an additional, more elaborate regulatory role for TOR signaling. Interestingly, a previous relationship between homoserine biosynthesis and Cdk8 activity was also noted in that accumulation of β-aspartate semialdehyde (ASA) caused by disruption of *hom6* (Figure S3) promotes both Cdk8 and Pho85-dependent degradation of Gcn4, but the mechanism for this effect has not been established (44).

Using a semi-intact cell assay for TORC1 (40), which we confirmed to be inhibited by rapamycin (Figure 6), we found that kinase activity recovered from *hom3* yeast was unaffected relative to WT. This may indicate that *hom3* mutations might affect TOR signaling downstream of TORC1, or that inhibition is abrogated during preparation of the assay samples. For example, it is interesting that *in vitro* TORC1 activity was stimulated by addition of glutamine or cysteine, but inhibited by aspartate (40). Consequently, it is possible that loss of Hom3 catalytic activity might cause elevation of intracellular aspartate that could inhibit TORC1; in this case it might be expected that elevated aspartate levels would become lost during preparation for the assay. To examine this possibility, we examined whether elevated aspartate in the media inhibited growth of *gal3* yeast on EB-gal but did not observe an effect (not shown); an obvious caveat is that higher media concentrations of aspartate may not necessarily produce elevated intracellular levels. Hom3 protein was also identified in complexes with Cdc55 in affinity capture MS proteomics studies (45), but the significance of this interaction is unknown.

The function of Cdk8 and the partially redundant Cdk19 protein kinase in humans are of considerable interest because a significant proportion of cancers may have alterations in their activity, particularly of melanoma and colorectal origin (1, 46). Cdk8/19 have enigmatic biological effects because phosphorylation of their target substrates have positive or negative effects on activity of specific factors and transcription of target genes (47). Currently, there are no known upstream regulators of Cdk8 kinase activity. The *gft* mutant screen was designed to address this issue, and characterization of one mutant revealed a role of TOR signaling for regulation of *GAL* gene expression. The long-term adaptation phenotype is among the earliest phenotypic differences described between various laboratory strains of *Saccharomyces cerevisiae*, and extensive analysis of this phenotype had indicated an important role for nutrient signaling regulating delayed *GAL* gene induction (13, 32, 48). Consequently, our discovery that TOR signaling modulates induction of *GAL* expression is consistent with these previous observations. Considering that these signaling molecules are conserved amongst eukaryotes, it will be interesting to determine whether mTORC1 activity may regulate Cdk8/19 target factors in human cells.

## Materials and Methods

### Yeast strains, plasmids, yeast techniques, and immunoblotting

All strains are derived from the W303-1A background and are detailed in Table 1, and DNA plasmids described in Table 2. Gene disruptions and reporter gene integrations were produced by homologous recombination using standard PCR-based methods and confirmed by PCR analysis of chromosomal DNA (49). Additional yeast genetic manipulations were performed as described (50). The screen for defects in Cdk8-dependent *GAL* induction was performed by u.v. mutagenesis of *gal3* W303 strain YJR7::131; colonies of mutagenized cells from minimal medium containing glycerol and lactic acid as the sole source of carbon were replica plated onto EB-gal plates. Mutants incapable of growth on EB-gal were transformed with a plasmid expressing WT *GAL3* and re-examined for growth on EB-gal; mutants incapable of growth when *GAL3*^+^ likely represented *gal* mutants and were discarded. Complementation analysis was performed with the remaining mutants; *gft* mutant strains were back-crossed to WT *gal3* W303 a minimum of 6 times. The Hom3 D297A mutation in plasmid pNH01 was created by site directed mutagenesis using oligos (GGCTATACCGcaTTATGTGCCG/ ACGACCAACACCATTCAG) with the NEB Q5 SDM system.

**Table 1:**
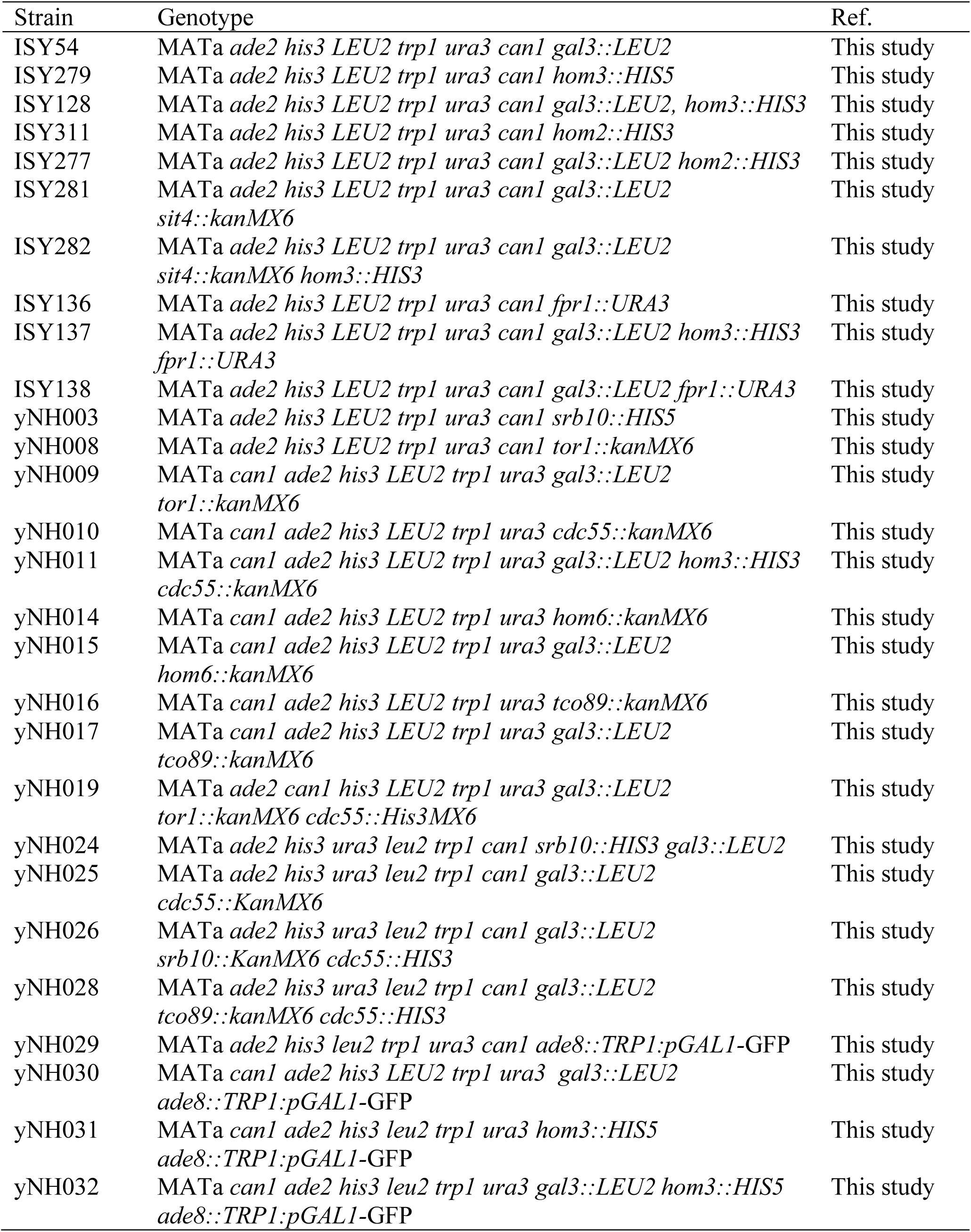
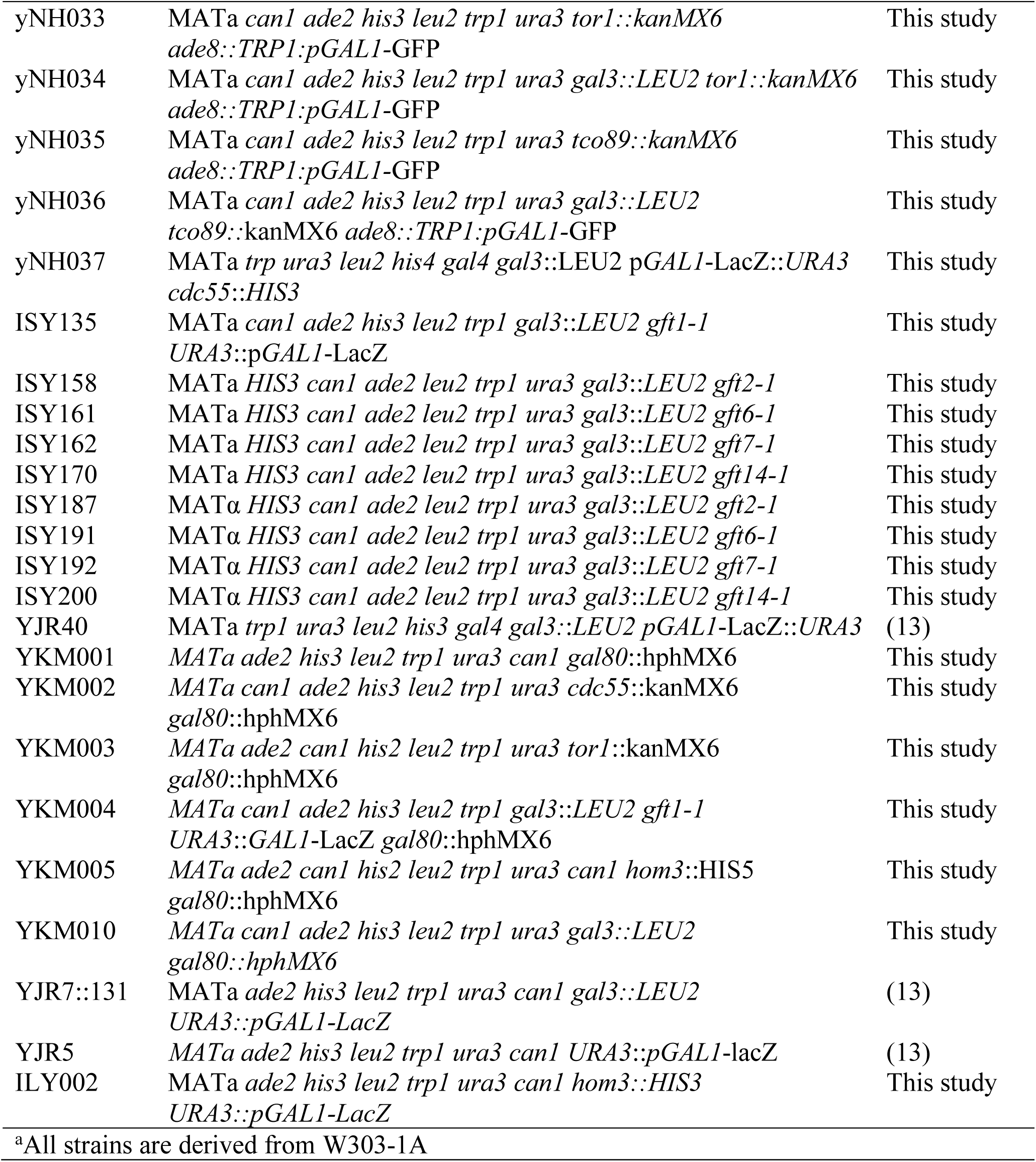
Yeast Strains^a^

**Table 2:** Plasmid DNAs

| Plasmid | Description | Ref. |
| --- | --- | --- |
| pRS314 | *TRP1*; ARS-CEN | (52) |
| pIS556 | *TRP1*; ARS-CEN; TEF1-Hom3-3X FLAG-6 | This study |
| pIS574 | *TRP1*; ARS-CEN; TEF1-Hom3 | This study |
| pNH01 | *TRP1*; ARS-CEN; TEF1-Hom3 (D297A) | This study |
| pIS297 | *TRP1*; ARS-CEN; *HOM3* | This study |
| pIS484 | *URA3*; ARS-CEN; TEF1-Srb10/ Cdk8-3X-FLAG-6his | This study |
| YCpG4trp | *TRP1*; ARS-CEN; *GAL4* | (16) |
| pRD038 | *TRP1*; ARS-CEN; *GAL4* (*S699A*) | (16) |
| pKW10Δ68<br>3 | *TRP1*; ARS-CEN; *GAL4*(Δ148–682) | (16) |
| pRS416-GAT-GFP | *URA3*; ARS-CEN; Gat1-GFP | (25) |
| pJP015 | *URA3*; ARS-CEN; TEF1-Srb10/ Cdk8-GFP | This study |
| pJR015 | *HIS3*; ARS-CEN; *GAL3* | This study |

Protein samples for immunoblotting were prepared by extraction with LiAc and NaOH (51). Antibodies against yeast aspartate kinase (Hom3) were produced in rabbits against a GST-Hom3 fusion protein expressed and purified from *E. coli*. Antibodies against the Gal4 DNA binding domain were produced in rabbits against 6-His-Gal4 (1-147) purified from *E. coli*.

### Flow cytometry, reporter gene quantitation and fluorescence microscopy

Strains with *GAL1*-GFP reporters were grown at 30°C in SC media containing glycerol, lactic acid and ethanol as the sole carbon source, to an O.D. A_600_ ∼ 1.0 and then induced with 2% galactose. Samples were taken at the indicated times and diluted in PBS to 400,000 cells per ml and analyzed for GFP expression and side scatter using a Guava Easycyte Flow Cytometer (MilliporeSigma). Mean fluorescence intensity was calculated using FlowJo software. Expression of the *GAL1*-LacZ reporter was determined by assay of β-galactosidase activity from permeabilized cells as previously described (50).

To monitor subcellular localization of Gat1, WT, *hom3* and *tor1* strains were transformed with plasmid pRS416GAT, expressing GAT1-GFP or pJP01, expressing CDK8-GFP as a nuclear marker. Cells were grown in SC lacking uracil at 30°C to an O.D. A_600_ ∼ 0.8. A 1 mL culture was mixed with 5 µL of (1/100 dilution) Hoeschst-3342 stock solution, and incubated for 30 minutes at 25 °C with gentle mixing. Cells were collected by centrifugation 2000 g for 4 minutes at RT, and 90% of the supernatant was removed. Cells were resuspended in the residual volume and 2 µL spotted onto a slide for analysis by microscopy.

### Protein kinase assays

TORC1 kinase assays *in vitro* were performed as described (40), but with several alterations. Cultures were grown to mid log phase in 50 mL YPD; cells were collected by centrifugation, washed once with ice cold dH_2_O containing 2 mM PMSF, and resuspended in 1 mL 0.1 M Tris-HCL (pH 9.4), 10 mM DTT, and incubated for 10 minutes on ice. The cells were collected by centrifugation, suspended in 1 mL spheroplasting buffer (0.7M Sorbitol, 10mM Tris-HCl [pH 7.5], 1mM DTT, 20mM NaN3, 0.1 mg/mL Zymolase), and incubated at RT for 30 minutes. Samples were then centrifuged at 1000xg for 2 min at 4°C. The pellets were washed with 1 mL ice cold sorbitol buffer (1M sorbitol, 150 mM K acetate, 5 mM Mg acetate, 20 mM HEPES-KOH [pH 6.6]), and resuspended in 350 µL sorbitol buffer. Spheroplasts were flash frozen in liquid nitrogen and stored at −80 °C in aliquots until use. In preparation for the assay, 20 μl spheroplast aliquots were thawed for 1 min at 30 °C and diluted with 1 mL of buffer A (0.25M sorbitol, 150 mM K acetate, 5 mM Mg acetate, 20 mM HEPES-KOH [pH6.6], 200 μg/mL PMSF). The samples were incubated on ice for 5 min and centrifuged at 13,000xg for 1 min, and the pellets resuspended in 1 mL buffer A. The pellets were washed a further time with buffer A, with buffer B (0.05 M sorbitol, 150 mM k acetate, 5 mM Mg acetate, 20 mM HEPES-KOH [pH6.6], and 200 ug/mL PMSF), and again with buffer A. The resulting semi-intact permeabilized cells were suspended in buffer C (0.4M sorbitol, 150 mM K acetate, 5mM Mg acetate, 20 mM HEPES-KOH [pH6.6]) at an O.D. A_600_ of 0.7. For each assay, 1 mL of the semi-intact cells were spun down at 13,000xg for 1 min and resuspended in 18 μL reaction buffer (0.4M sorbitol, 150 mM K acetate, 5mM Mg acetate, 20 mM HEPES-KOH [pH 6.6], 40 mM creatine phosphate, 200 ng/ul creatine kinase, 1 mM Pefabloc SC, 4 μg/mL aprotinin, 1 μg/mL pepstatinA, 2 μg/mL leupeptin, 0.3 μg/mL 4EBP1, Sigma Aldrich). Rapamycin was added at 1 or 10 µM. Reactions were initiated by addition of 2 µL of ATP-amino acid mix (5 mM ATP with 2 % amino acid mix) and incubated for 10 minutes at 30°C. Reactions were terminated by addition of 2X SDS PAGE sample buffer, and analyzed by SDS-PAGE and immunoblotting with anti-4EBP1 antibodies and anti-Phospho-4EBP1 (Cell Signaling).

To measure Cdk8 kinase activity *in vitro*, yeast with the indicated genotype expressing Srb10/ Cdk8-3XFLAG were grown in SD-Ura to an O.D. A600 ∼ 0.8 and recovered by centrifugation. The cells were washed twice in kinase lysis buffer (KLB) (50 mM Tris [pH7.5], 5mM ETDA, 200mM NaCl, 0.1% NP-40, and protease inhibitors), and resuspended in 1 ml KLB per ml culture. Cells were lysed by vortexing with glass beads, and the lysates clarified by centrifugation at 13,000Xg for 10 minutes at 4°C. FLAG-tagged Srb10/ Cdk8 was recovered by immunoprecipitation with anti-FLAG-conjugated M2 agarose beads (Sigma). Samples were washed two times in KLB followed by two times in kinase assay buffer (KAB) (10 mM MgCl2, 50 mM Tris [pH 7.5], 1 mM DTT and protease inhibitors), and the beads suspended in 50 µL KAB. Reactions contained 5 µL of the bead suspension and 1 µg substrate protein in 10 µL KAB, and started by adding 2 pmol [γ-32P]ATP, and were incubated at 30 °C for 20 minutes. Reactions were stopped by addition of 10 µL 2X SDS-PAGE sample buffer; the samples were boiled for 2 min, and centrifuged 13,000Xg for 1 min. The supernatant was resolved on 10% SDS PAGE, and the gel dried and exposed to Kodak Biomax film. GST, and GST-CTD substrate protein were expressed and purified from *E. coli* (15), and WT Gal4 from insect cells using baculovirus (12).

## Acknowledgements

We thank LeAnn Howe and Maria Aristizabal for helpful comments. This research was supported by funds from the Natural Sciences and Engineering Research Council (F12-04577).

## Legends to Supplementary Figures

**Figure S1:**
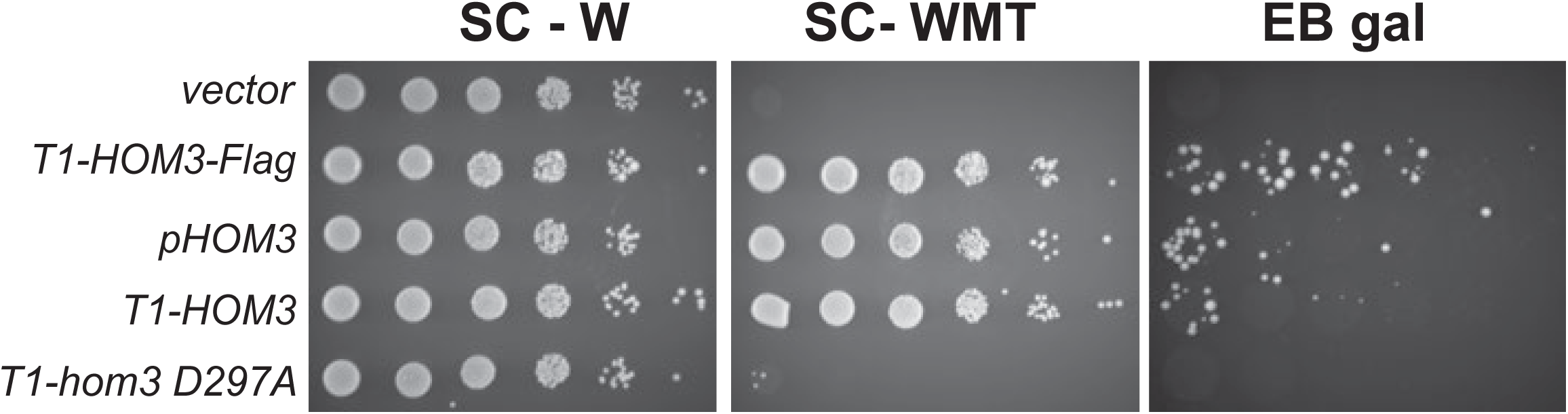
Plasmids expressing *HOM3* complement defective growth of *gft1-1 gal3* yeast on EB-gal. Yeast strain ISY135, bearing the *gft1-1* mutation was transformed with plasmids, including a vector control (pRS314), expressing Hom3-3XFlag expressed from the *TEF1* promoter (pIS556, T1-*HOM3-Flag*), bearing a genomic clone of *HOM3*, recovered in complementation analysis of the mutant phenotype (pIS297, *pHOM3*), or vectors expressing the WT (pIS574, *T1-HOM3*) or D297A mutant (pNH01, *T1-hom3* D297A) ORF from the *TEF1* promoter. Cultures were diluted and spotted onto SC lacking tryptophan (SC-W), SC lacking tryptophan, methionine and threonine (SC-WMT), or EB-gal, from 10-fold serial dilutions.

**Figure S2:**
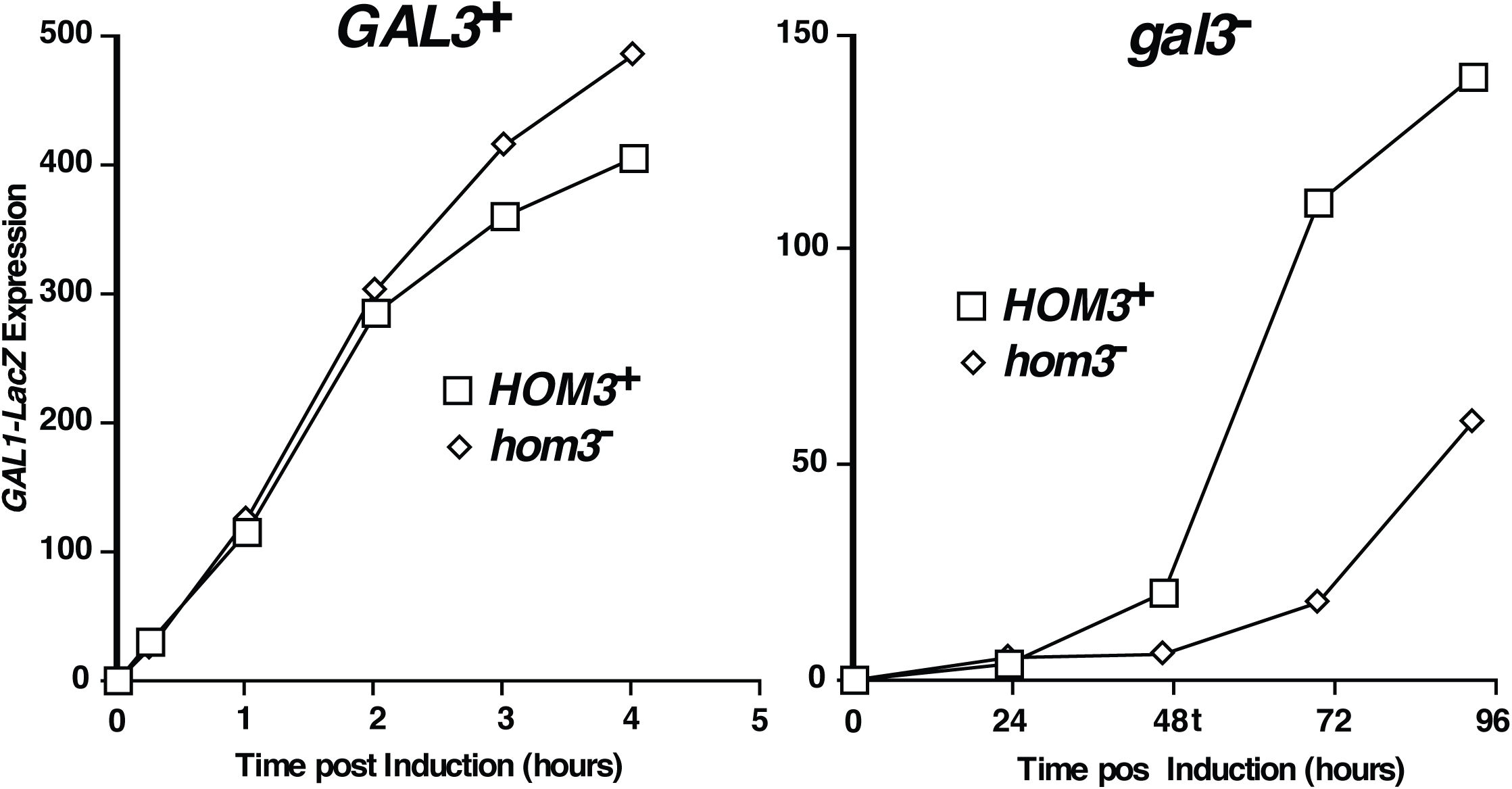
Mutation of *hom3* inhibits induction of *GAL1* expression in *gal3* yeast. W303-1A (WT, left) and (*gal3*, right) yeast expressing WT HOM3 (□) or a *hom3* deletion (⋄) and bearing a *GAL1*-LacZ reporter were induced with 2% galactose for the indicated times (hrs), and assayed for β-galactosidase activity.

**Figure S3:**
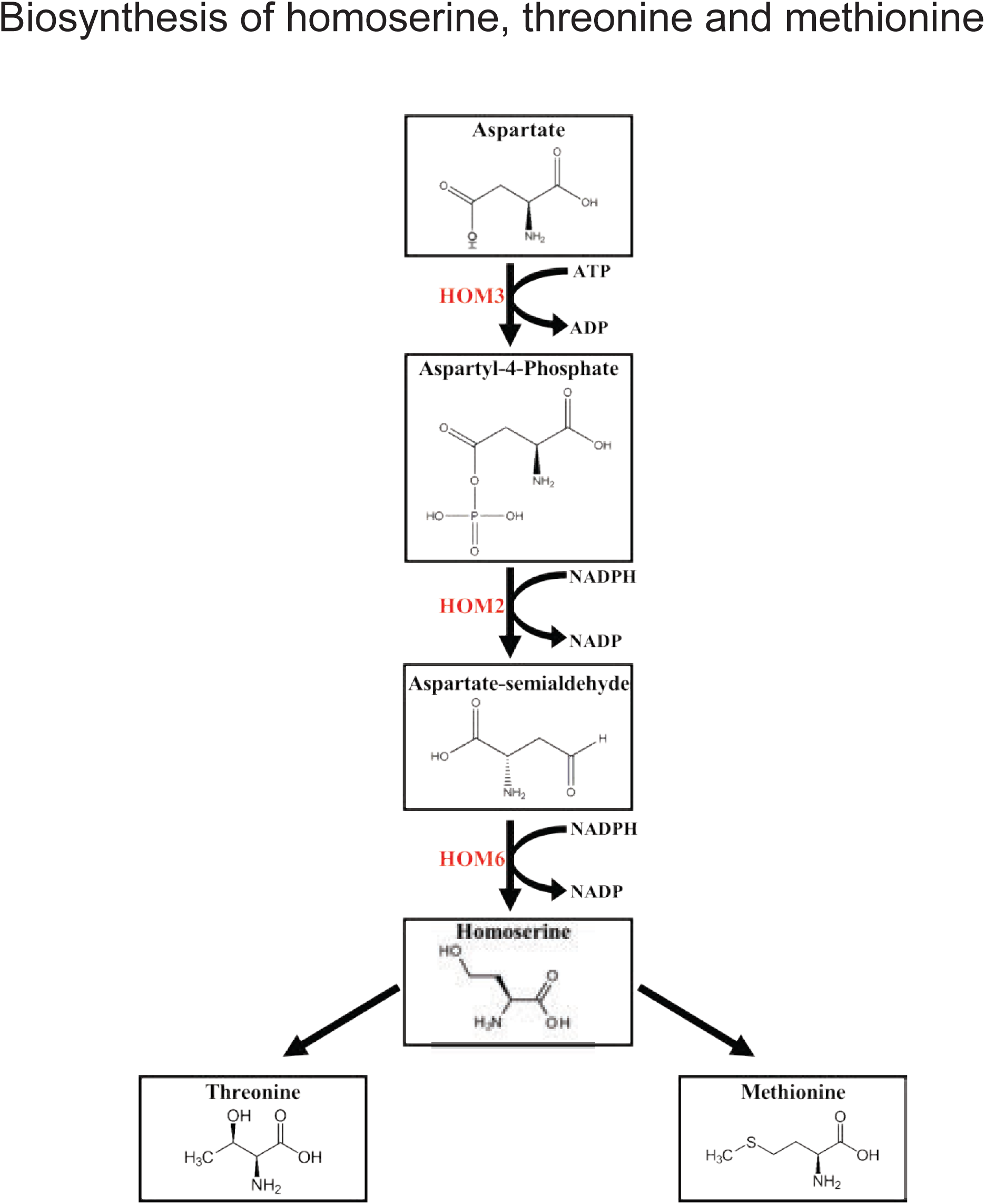
Biosynthetic pathway in yeast for homoserine, threonine and methionine. Genes encoding enzymes for biosynthesis of homoserine include *HOM3* (aspartate kinase), *HOM2* (aspartic beta semi-aldehyde dehydrogenase) and *HOM6* (homoserine dehydrogenase).

**Figure S4:**
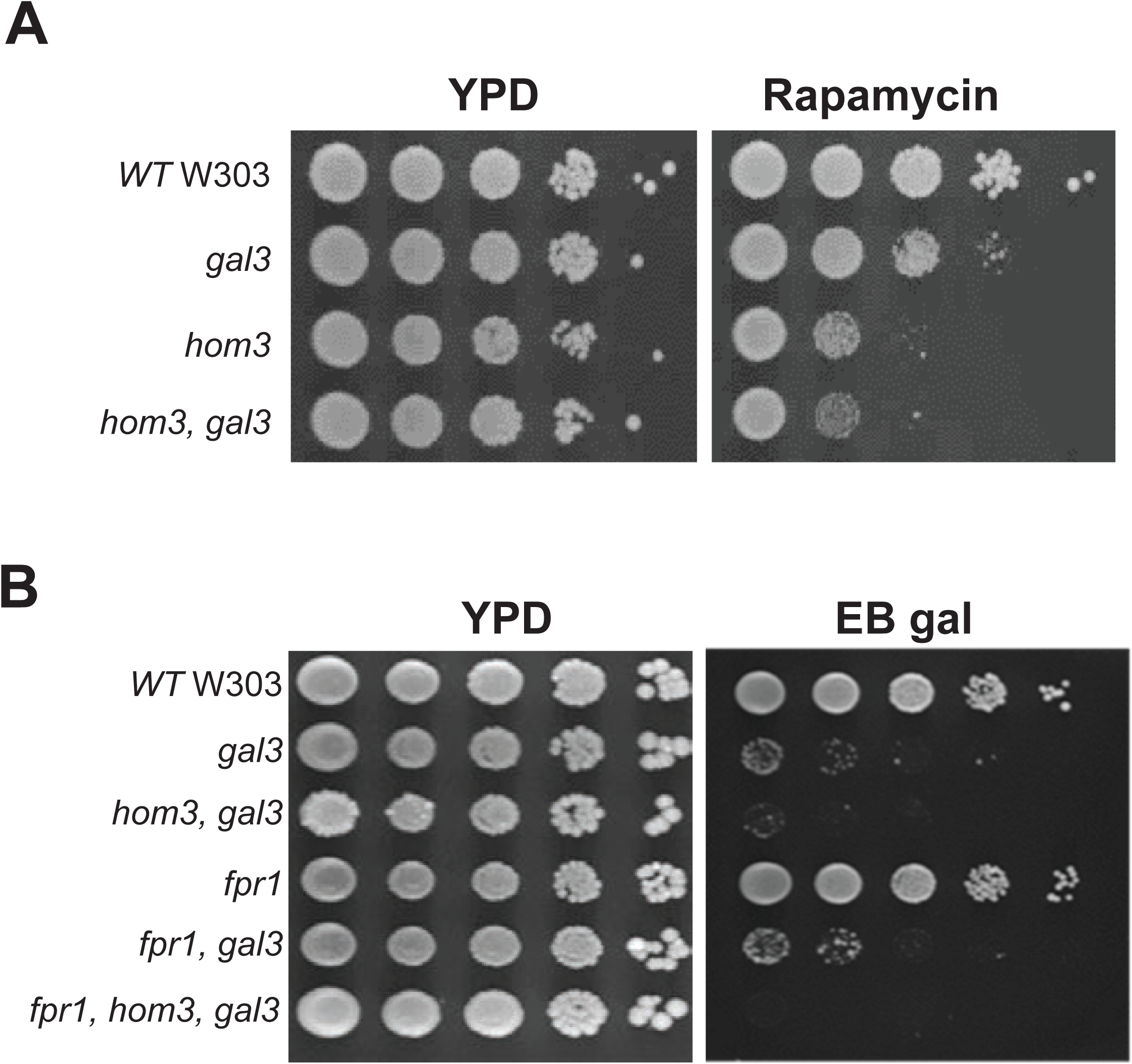
Disruption of *hom3* causes sensitivity to sub-lethal concentrations of rapamycin. **Panel A:** Strains with the indicated genotype were spotted from overnight YPD cultures in 10-fold serial dilutions YPD or YPD containing 5 ng/mL rapamycin plates and incubated at 30°C for 3 days. **Panel B:** Strains with the indicated genotype were grown overnight in YPD, diluted to an O.D. A_600_ of 1.0, spotted onto YPD or EB-gal plates in 10-fold serial dilutions, and grown at 30°C for 5 days.

**Figure S5:**
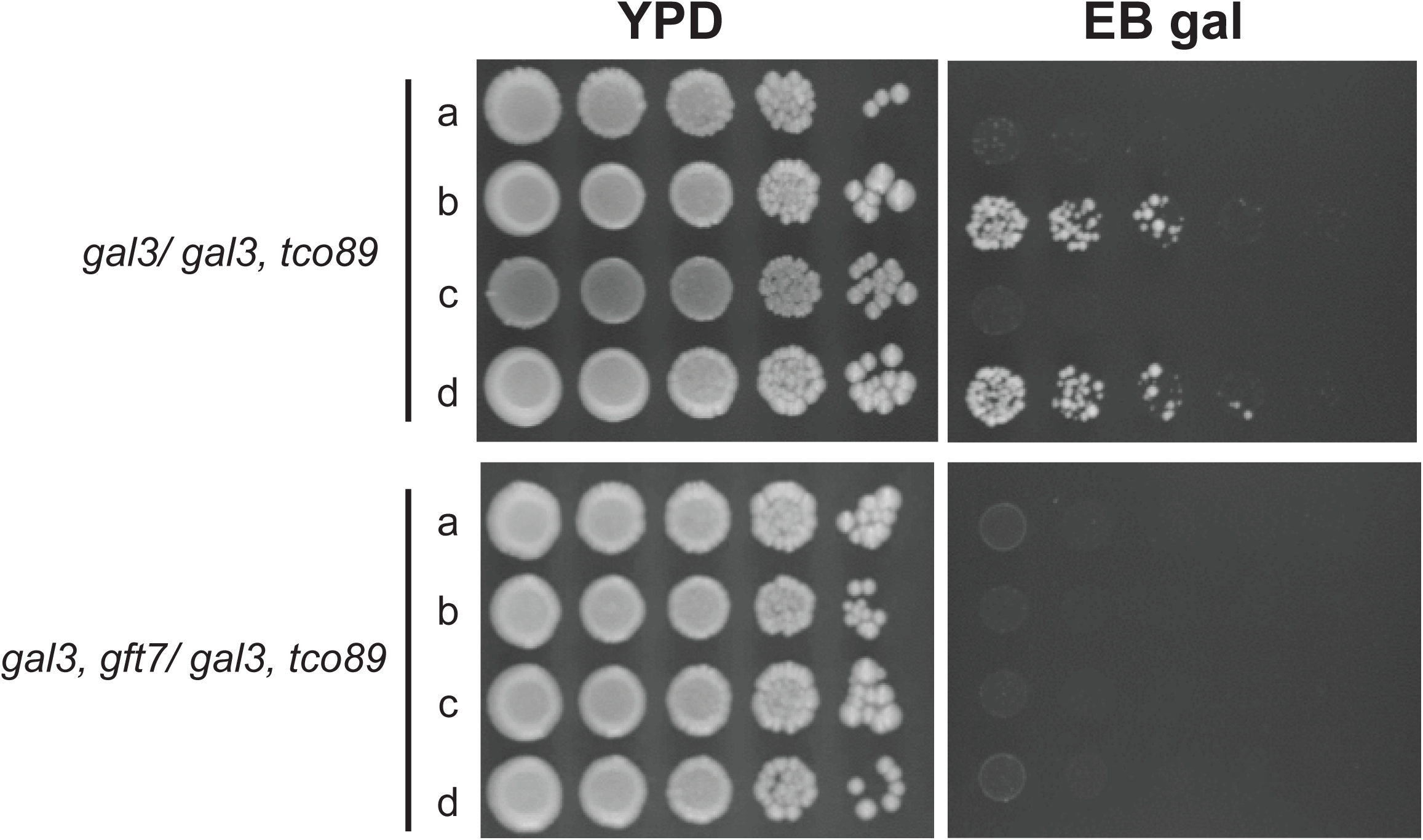
The *gft7-1* mutation is allelic to *tco89*. Diploid strains were produced by mating *gal3 tco89* (yNH017) with *gal3* (ISY54) (top panel), or *gal3 gft7-1* (ISY192) (bottom panel). Following sporulation, tetrads were dissected and haploids from spores were spotted from YPD cultures at 10-fold serial dilutions onto YPD or EB-gal plates. The plates were incubated for 5 days at 30°C.

**Figure S6:**
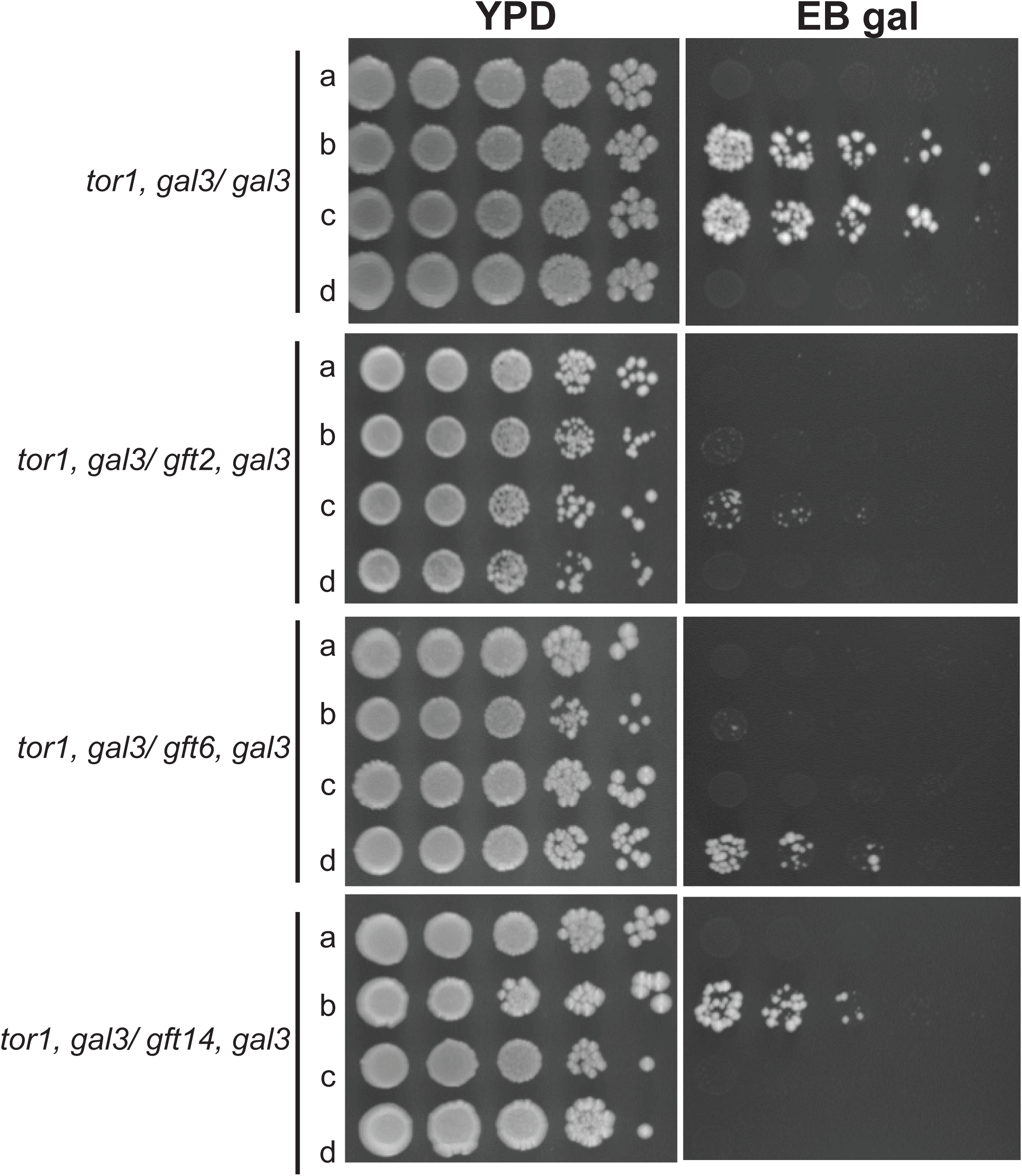
A *tor1* disruption produces unlinked non-complementation with *gft2-1, gft6-1* and *gft14-1.* Diploid strains were produced by mating *gal3 tor1* (yNH009) with *gal3* (ISY54), *gft2-1* (ISY187), *gft6-1* (ISY191), or *gft14-1* (ISY200). Tetrads were dissected from sporulated cultures and haploids from spores spotted onto YPD or EB-gal plates. The plates were incubated for 5 days at 30°C.

**Figure S7:**
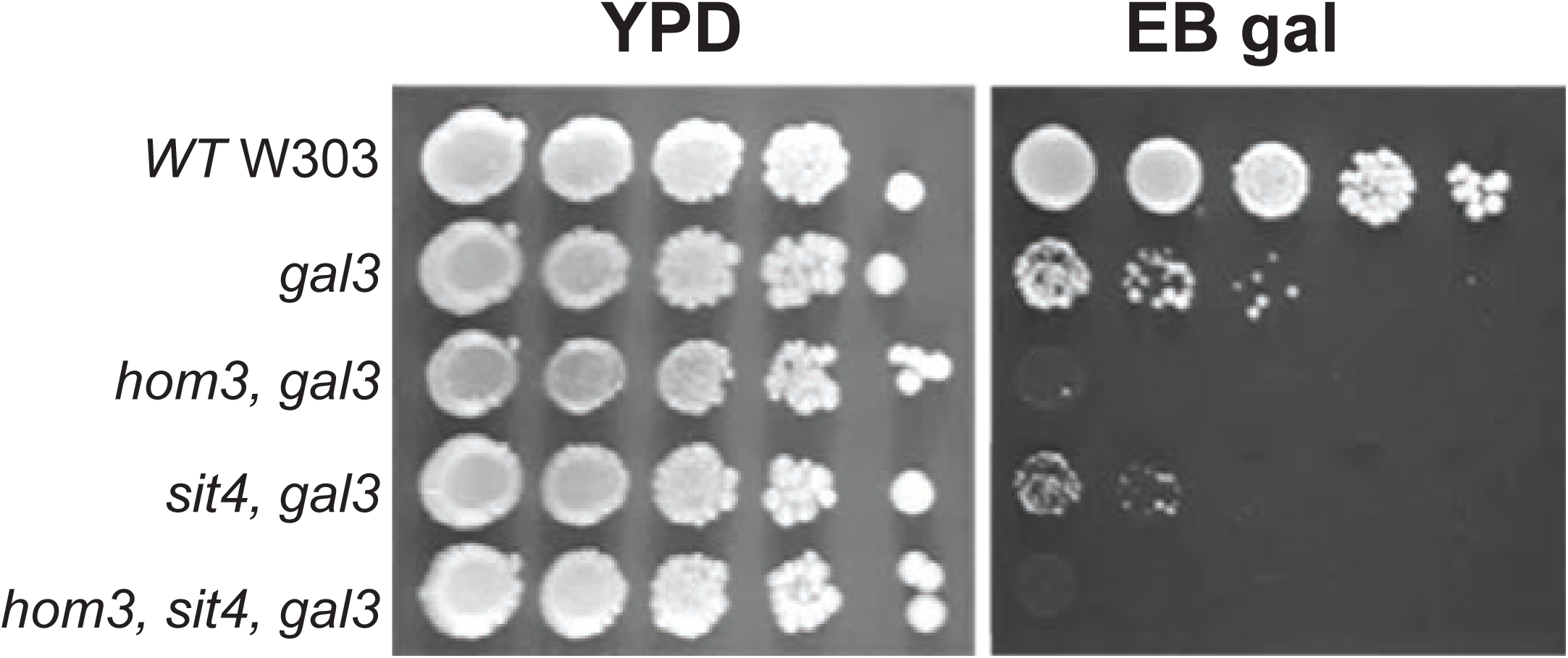
*GAL* expression in *gal3* yeast is unaffected by disruption of *sit4*. Strains with the indicated genotype were grown overnight in YPD, diluted to an O.D. A_600_ of 1.0, spotted onto YPD or EB-gal plates in 10-fold serial dilutions, and grown at 30°C for 5 days.

